# ABRIDGE: An ultra-compression software for SAM alignment files

**DOI:** 10.1101/2022.01.04.474935

**Authors:** Sagnik Banerjee, Carson Andorf

## Abstract

Advancement in technology has enabled sequencing machines to produce vast amounts of genetic data, causing an increase in storage demands. Most genomic software utilizes read alignments for several purposes including transcriptome assembly and gene count estimation. Herein we present, ABRIDGE, a state-of-the-art compressor for SAM alignment files offering users both lossless and lossy compression options. This reference-based file compressor achieves the best compression ratio among all compression software ensuring lower space demand and faster file transmission. Central to the software is a novel algorithm that retains non-redundant information. This new approach has allowed ABRIDGE to achieve a compression 16% higher than the second-best compressor for RNA-Seq reads and over 35% for DNA-Seq reads. ABRIDGE also offers users the option to randomly access location without having to decompress the entire file. ABRIDGE is distributed under MIT license and can be obtained from GitHub (https://github.com/sagnikbanerjee15/Abridge) and docker hub. We anticipate that the user community will adopt ABRIDGE within their existing pipeline encouraging further research in this domain.

## INTRODUCTION

Next generation sequencing (NGS) has opened up opportunities to study several biosystems from a quantitative viewpoint ((1, 2, 3, 4)). Over the years, numerous sequencing protocols have been designed to probe the modus operandi of number of biological processes ((5, 6)). Researchers have perfected these protocols - making them more economical and effective. This made sequencing accessible to even underfunded labs leading to a surge in data. Short read data (generated typically on Illumina platforms) is often mapped to a reference (genomic/transcriptomic) and then used for several purposes – assembling ((7, 8, 9, 10)), annotating ((11, 12, 13, 14)), finding differentially expressed genes ((15, 16)) and proteomics ((3, 17, 18, 19)). Most bioinformatics projects utilize a very large set of RNA-Seq or DNA-Seq samples collected from multiple tissue types and conditions. The primary step in such experiments is to align the RNA-Seq samples to a reference that generates a sequence alignment map (SAM) ((20)) that is stored in either a binary alignment map (BAM) ((20)) or compressed alignment file (CRAM) ((21)) format. Even though these formats offer compression to some extent, the total size of all the aligned files can often exceed the storage capacity that small labs can afford. Hence, better compression techniques are needed that utilize the underlying structure of reference alignment files and offer a multitude of options to cater to a diverse range of user requirements.

Short reads, generated by sequencing platforms like Illumina, need to be mapped to a reference using aligners like STAR ((22, 23)), HiSAT2 ((24)) or BWA ((25)) before further processing. These aligners typically output the result in a SAM format which can be converted to a binary BAM format to achieve better compression. SAM format stores the location, shape (CIGAR string) (https://genome.sph.umich.edu/wiki/SAM), nucleotide bases, quality scores and tag level information for each aligned read. Since alignments in SAM format are stored for each read, the file size grows linearly with the number of reads in the sample. Hence, there is a need to devise an algorithm that can exploit the underlying structure of SAM files and offer the best possible compression in a reasonable amount of time.

A considerable amount of time and effort has been directed to designing algorithms to compress alignment files to reduce storage demands and facilitate file transfers ((26, 27, 28)). Most approaches achieve compression by eliminating redundant data by accumulating alignment information across multiple reads or alignments. SAM compressors, like NGC ((29)), DeeZ ((30)) and genozip ((31)) are reference based while BAM, CRAM, Quip ((32)) and CSAM ((33)) are reference free. Reference-based approaches achieve compression by representing an aligned read with a description of how it differs from the reference. This eliminates the need to store the actual read sequence thereby reducing storage demands. Quality scores do not map to any reference and hence cannot be compressed like the read string. Hence some compressors like NGC, CSAM, genozip and DeeZ offers users the option to map quality values within a range to a single value. While this can lead to better compression, it might remove quality scores of mismatched bases which are essential for detecting single nucleotide polymorphisms (SNPs). Quip implements Markov chains to encode read sequences and quality scores. Samcomp ((34)) compresses SAM alignments in lossless fashion by tokenizing the read identifiers and sorting the reads as a reference difference model. A very similar approach is undertaken by DeeZ where tokenized read names and read sequence are compressed with delta encoding.

To overcome the shortcoming of previous SAM compression approaches, we introduce ABRIDGE. We offer users a plethora of choices to compress SAM files. To optimize space utilization, ABRIDGE accumulates all reads that are mapped onto the same nucleotide on a reference. ABRIDGE modifies the traditional CIGAR string to store soft-clips, mismatches, insertions, deletions, and quality scores thereby removing the need to store the MD string. To further reduce space demand, ABRIDGE modifies the CIGAR information to store the strand on which the read was mapped. ABRIDGE also offers the option to alter quality scores of nucleotide bases that had a perfect match with the reference thereby reducing even more space (**Supplementary Figure 7**). All features of multi-mapped reads are stored with their individual CIGAR strings. Hence reads mapping to homeologs in polyploid species will retain their alignment profile. Users can choose from three levels of compression offering varying extents of compression with the caveat of the duration of compressing. ABRIDGE offers options of completely lossless compression and selectively lossy conversions. Consequently, decompressions in ABRIDGE can regenerate the entire SAM file with or without modifications depending on the choices made during compression. In this manuscript, we explore the different modes in which ABRIDGE can operate and compare it with other state-of-the-art tools.

## MATERIALS AND METHODS

ABRIDGE accepts a single SAM file as input and returns a compressed file that occupies less space than its BAM or CRAM counterpart. Users can choose to retain all the quality scores which would initiate a lossless compression. In several applications, storing the entire quality score is redundant. Hence, ABRIDGE can be configured to preserve only those quality values which for which the corresponding nucleotide base was a mismatch to the reference or an insertion into the read sequence. This option considerably reduces the compressed size but stores the most relevant information which can later be used for analysis that uses quality scores (e.g. variant calling). To further reduce space, users can eliminate quality scores altogether. Some downstream software like transcriptome aligners do not use soft-clips or mismatches, so we designed ABRIDGE to provide options to ignore such information in the SAM file while compressing. ABRIDGE compresses SAM files in two passes – in the first pass, relevant information from the SAM file is rearranged and in the second pass, the file is compressed using generic compressors. ABRIDGE decompresses data by applying the reverse algorithm producing all the requested information to be stored during compression. Once the data is compressed, users can retrieve alignment information from random locations making it very easy to access alignments from anywhere in the genome without having to decompress the entire file.

ABRIDGE achieves a high compression ratio due to the underlying strategies of eliminating redundant data. Instead of storing the entire sequence of reads, ABRIDGE stores the location of the reference to which the read mapped and relevant information about the mismatched and/or inserted base pairs. Instead of storing the exact mapped location, it keeps the difference in mapped position from the previous alignment. This saves a substantial amount of space for both RNA-Seq and DNA-Seq data. ABRIDGE also merges the exact same reads originating from the same nucleotide position of the reference. Read names for uniquely mapped single-ended reads are discarded but are preserved for multi-mapped single ended reads and for paired-ended reads to associate each read with the corresponding fragment. ABRIDGE offers users a multitude of choices for storing quality values. Users can request to store all the quality values without making any changes or allow ABRIDGE to modify the quality scores of some bases to facilitate better and faster compression. Instead of blindly modifying the quality scores, ABRIDGE inspects each base pair and modifies its quality value only if the base pair was aligned perfectly to the reference. Hence, the quality scores of bases which are inserts and/or mismatches are preserved. This provides the users with the opportunity to retain all the relevant information necessary to perform vital downstream analysis. ABRIDGE stores a modified version of the CIGAR string by including soft clipped bases, quality scores of mismatched and inserted bases along with nucleotides that did not match with the reference. Users are also provided with the choice of achieving best compression by eliminating quality scores altogether. This option is helpful for storing alignments files for the purpose of performing transcriptome assemblies where quality scores are not typically used ((10)).

Unlike the read sequence, quality scores cannot be “mapped” to any reference. Hence ABRIDGE stores quality values as reported and then compresses those with generic compressors. ABRIDGE can store quality values in four different ways – (1) Discard quality values of reference matched bases and include only the mismatched and inserted bases. For this case, quality values are stored within the enhanced CIGAR, (2) Store all quality scores with altered values for reference matched bases, (3) Store all quality values without making any change in the quality values, and (4) Discard quality scores altogether.

Information about the alignment of each read is typically stored in the CIGAR and the MD string. While CIGAR string can indicate the soft-clips, matches, insertion and deletions, it is not designed to store mismatched nucleotides and read inserts. MD string, on the other hand, reports the mismatched bases. Hence, both the CIGAR string and MD string are needed to accurately reconstruct the appropriate alignment of the read to the reference. Since the CIGAR string and the MD string contain overlapping information we decided to integrate them and generate a single representation which we call the integrated CIGAR. The integrated CIGAR contains complete information from which the entire alignment can be reconstructed (**Figure 1**). Quality scores are stored within the integrated CIGAR if the user requests for it. Quality scores for only the mismatched bases and the inserts are stored. If the user requests to store quality scores for all the nucleotide bases then the scores will be stored in a separate file. An illustration of how the integrated CIGAR string is constructed has been provided in **Figure 1**. Once each alignment entry is encoded, an index file is generated that can speed up file access in the future. The index contains information about the location of a pile of reads. During random access, the entire index file is read into the memory. According to the request made by the user, a specific portion of the compressed file is read and subsequently decompressed. Consulting the index file eliminates the need to decompress the whole alignment thereby speeding up random access. Finally, Generic compressors are used to compress the index along with the concise alignment file.

**Figure 1.**
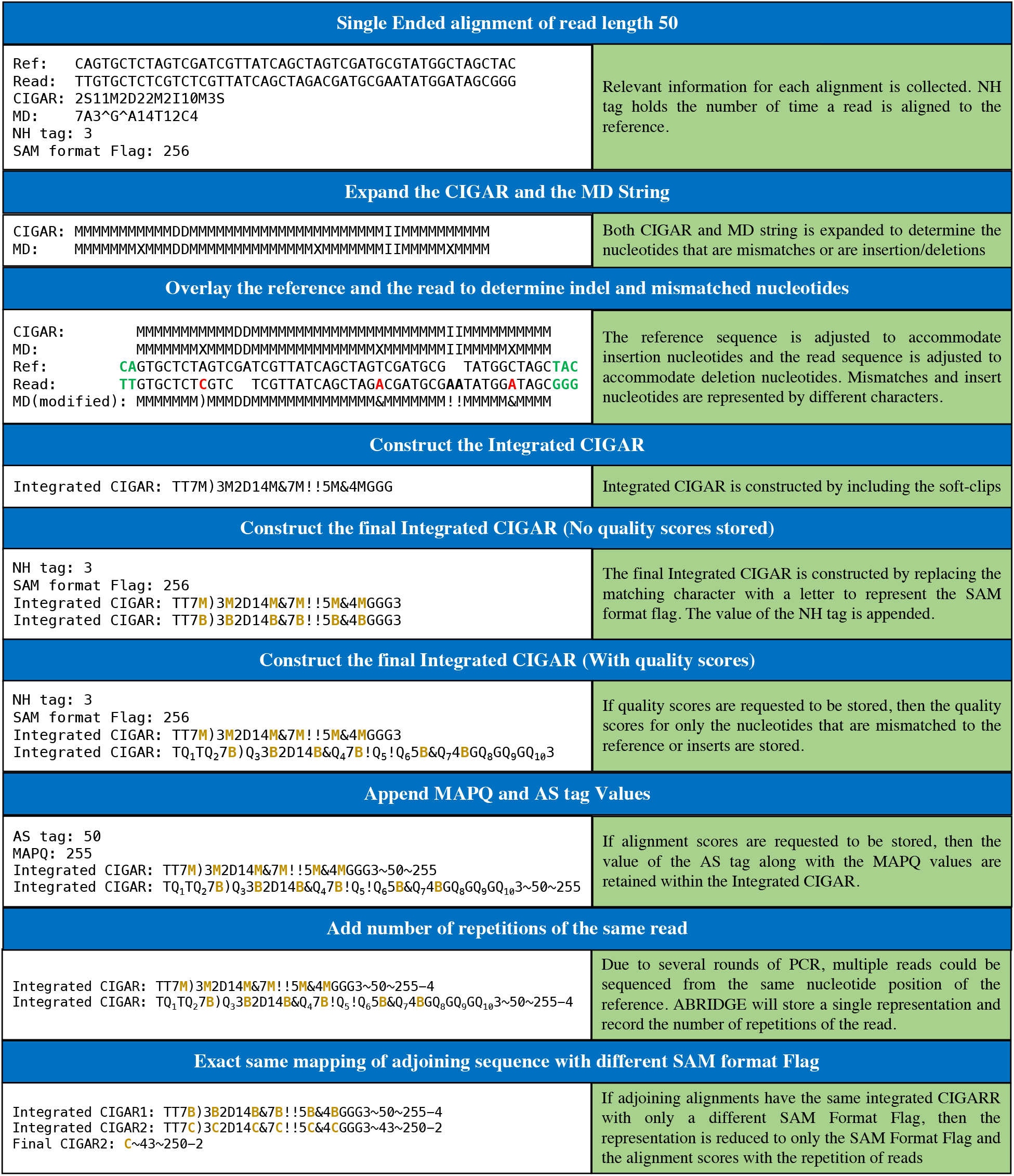
Generation of integrated CIGAR. Each alignment is converted to a string that combines both CIGAR and the MD. The CIGAR and the MD string is expanded to determine the locations of insertion, deletions and mismatches. Soft clips are removed from the front and the end of the read. The reference and the read sequence is consulted to locate the mismatches. The insert nucleotides and the mismatch nucleotides are replaced with special characters. Quality scores are also inserted into the integrated CIGAR for insert and mismatched nucleotides

ABRIDGE will generate the compressed file in ‘.abridge’ format which is essentially several files compressed using one of the three compressors - Brotli, 7z or ZPAQ. During decompression, a SAM is produced from the compressed files. The decompression step might require substituting dummy quality scores for some cases, depending on how the quality scores were stored during compression. The decompressed file will be sorted and dummy read names will be produced where they were discarded to save space. Some applications, like genome-guided assembling, do not require the nucleotide sequence. Hence, ABRIDGE allows the user to decompress without generating the actual read sequence. This option is faster to execute since it does not require the reference to be used.

## RESULTS

We tested ABRIDGE on RNA-Seq and DNA-Seq data of various read depths (**Supplementary Table 2**). Programs were executed on a cluster with Intel(R) Xeon(R) CPU E5-2670 v2 processors with 2.50 GHz. CentOS Linux release 7.9.2009 was the operating system. ABRIDGE is entirely written in C and gcc compiler (v4.8.5) was used. We carried out experiments using different parameter settings as described in **Table 7**. The first parameter setting produces lossless compression and then we demonstrate how ABRIDGE can be configured to retain the requested information without impacting downstream applications.

Details about data acquisition and processing have been mentioned in **Supplementary document**.

### SAM file format requirements

Input to ABRIDGE, and also to the other programs, need to be provided in SAM format. The file must be sorted by position and should have a proper SAM header. In addition, each alignment must have three tags - NH, MD, and XS. NH tag stores the number of times the read has been mapped which assists ABRIDGE to distinguish between uniquely mapped and multi mapped reads. MD tag contains information about mismatched bases and deletions which are used to generate a field in the compressed file. XS tag stores information about the strand to which the read was aligned.

### ABRIDGE achieves the best lossless compression

ABRIDGE has two major goals - (1) achieve a high level of lossless compression, and (2) provide users with different modes of compression. Lossless compression is achieved by preserving only non-redundant information from SAM alignment file. Alignment files in SAM format were provided as input to the compression software. ABRIDGE performs the best compression owing to the usage of zpaq compressor (**Table 1**). For single-ended reads ABRIDGE discards the read names for uniquely mapped reads. But for paired end reads, ABRIDGE needs to store the read names of both the pairs to enable associating the reads with the same fragment during decompression. This causes a slightly poor compression performance of ABRIDGE (with 7z) (**Figure 2**). CSAM generates a file which is larger than the CRAM file itself. SAMCOMP attains the second best compression for paired-ended reads and third best for single ended reads. GENOZIP and DEEZ exhibit average performance in terms of ratio of compression.

**Table 1.**
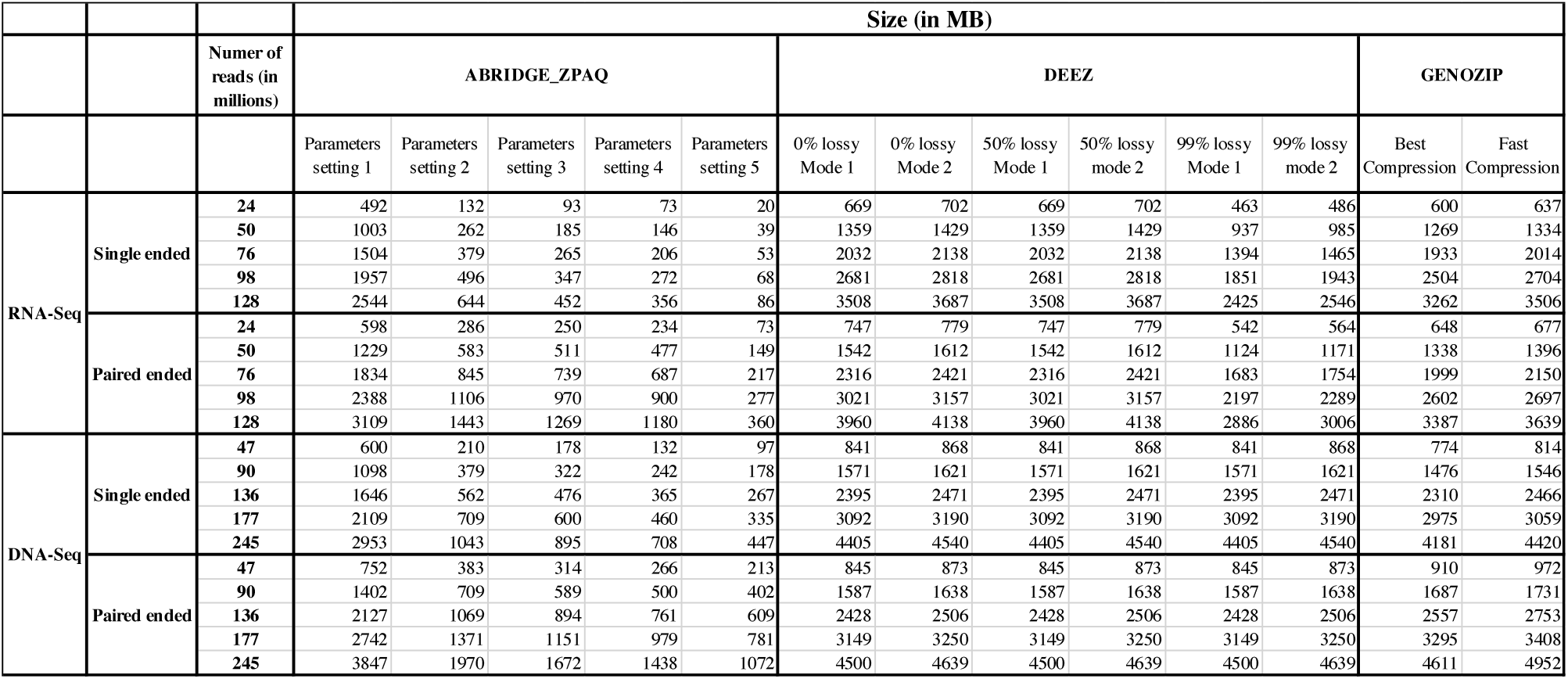
Comparison of lossless and lossy compression with different software.

**Figure 2.**
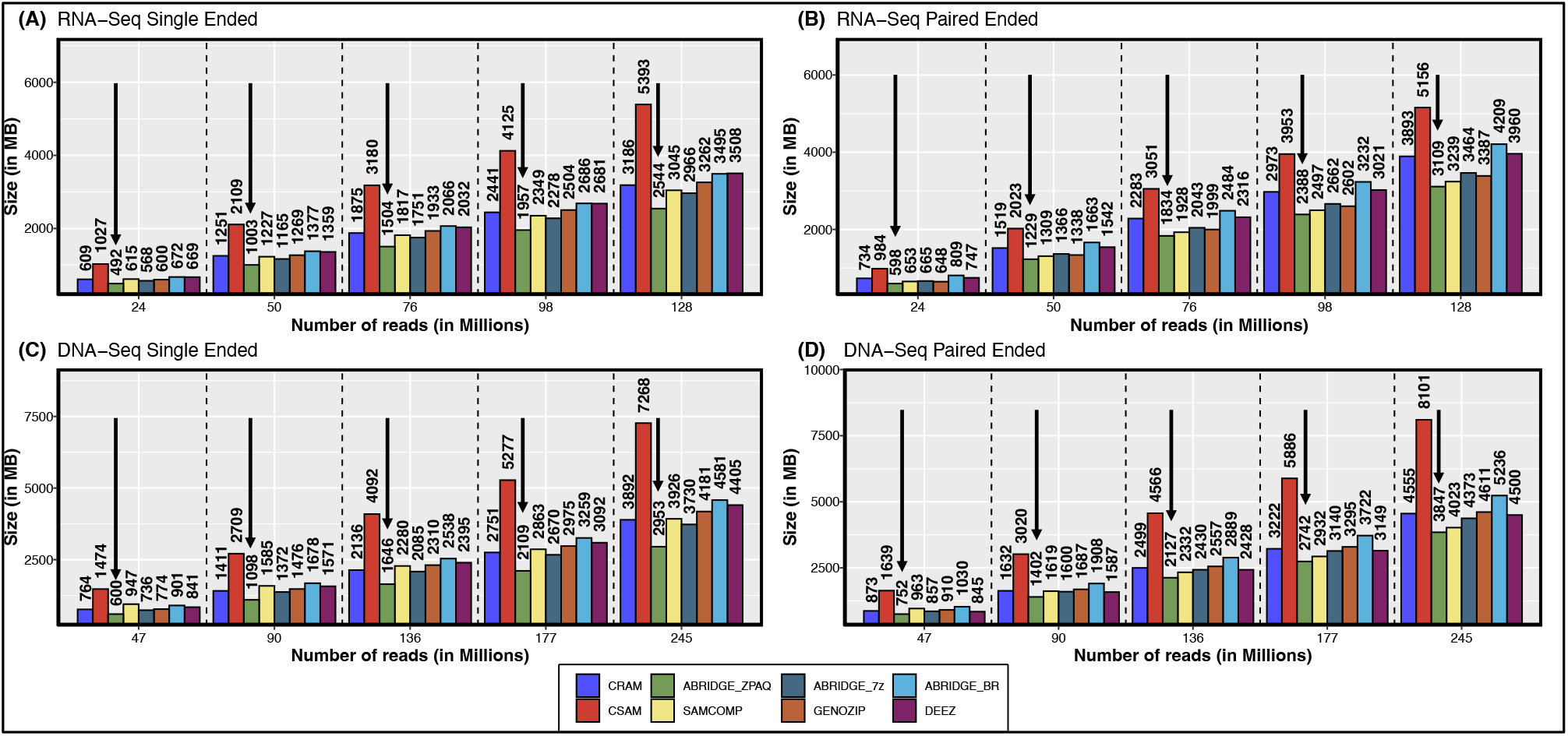
Compression achieved by different software. Compressed sizes of files produced by each compression software for RNA-Seq, DNA-Seq, Single ended and Paired ended data. Only software that are regularly updated and maintained have been included in the analysis. Each segment represent compression for increasing file sizes.

### ABRIDGE offers multitude of options for Lossy compression

ABRIDGE offers users with different options of compression as outlined in **Table 7**. Instead of blindly compressing quality scores, ABRIDGE offers users to modify quality scores of those nucleotide bases that perfectly match with the reference. This allows the user to retain the exact quality score of mismatched bases and insertions which would be useful for downstream analysis. With parameter setting number 2, ABRIDGE converts the quality score of matched bases to facilitate vertical run-length encoding leading to higher compression resulting in lower file size (**Supplementary Table 3**). This is further illustrated in **Supplementary Figure 6** where the space requirement for storing quality scores greatly reduces from parameter setting 1 to parameter setting 2. The next set of parameters discard all quality scores except for the non-matched bases. The complete discard of quality scores leads to a further reduction in space requirements. The fourth parameter setting removes all quality scores, soft-clips, and mismatched bases. Since these did not occupy too much space, their removal did not reduce space significantly. In the final parameter setting, only the position of the mapped reads are preserved leading to the smallest file size. As expected, ZPAQ produces the best compression followed by 7z (**Supplementary Table 3**).

Other software also offer the provision of lossy compression. Both DEEZ and GENOZIP were executed with different parameter settings of lossy compression. ABRIDGE lossy compression, with approximates quality scores (parameter setting 2) was able to produce a better compression than all other software operating in lossy mode (**Table 1**).

### ABRIDGE quickly compresses data

We compared the duration required to compress the SAM files. Even though ABRIDGE was not able to compress data the fastest, it was comparatively faster than CSAM and GENOZIP **Supplementary Figure 4**. The main bulk of time is taken by the generic compressors (brotli, 7z and ZPAQ) which can be improved by allocating more CPU cores. Compression of a file is performed only once, hence we believe users will not be hesitant to dedicate the time. **Supplementary Table 4** lists the duration of compression for the three generic compressors used in ABRIDGE along with different modes of compression. The duration of compression reduces with more lossy compression for both Brotli and 7Z. It is interesting to note that for ZPAQ the duration does not change much.

### ABRIDGE decompresses data faster than other software

In order to utilize the alignments, the compressed files by ABRIDGE need to be decompressed. Unlike compression, decompression needs to be done multiple times depending on how often the alignment files are required to be accessed. Hence, we offer users the choice of multiple compressors that can help decompress files quicker. As depicted in **Supplementary Figure 5**, 7z can decompress files very quickly. Unfortunately, ZPAQ takes the most time to decompress files even when it offers the best compression. Both brotli and 7z take almost the same time to decompress files which were compressed using different parameter settings (**Supplementary Table 4**). ZPAQ, on the other hand, decompresses files faster when the compression was lossy.

### ABRIDGE can retrieve data randomly from any location

During compression, ABRIDGE creates an index to facilitate random search. We compared the duration of generating the ABRIDGE index with the time taken to generate the BAM or CRAM index. As listed in **Supplementary Table 6**, CRAM takes the least time to generate indices. BAM and ABRIDGE take almost the same amount of time for single-ended reads. ABRIDGE takes longer for paired-ended reads since it needs to navigate through all the read names to index the file. ABRIDGE consume more memory to generate the indices whereas BAM and CRAM consumes much less memory. Interestingly, CRAM takes the same amount of memory for generating index even when the number of alignments increase.

Random access with ABRIDGE involves decompressing the file and then randomly accessing the requested location. Since ABRIDGE decompresses the entire file, it takes much longer to access random locations than CSAM, GENOZIP, BAM and CRAM. DEEZ takes the longest to access locations randomly since it decompresses the entire file (**Supplementary Table 9**). Both BAM and CRAM consume the least memory while ABRIDGE consumes the most (**Supplementary Table 8**)).

## DISCUSSION

We present ABRIDGE - a state-of-the-art software for compressing SAM alignments. ABRIDGE compresses alignments after retaining only non-redundant information. It achieves superior compression by merging similar reads mapped to the same location of the reference and encoding only those nucleotides that deviate from the provided reference. Strand information is also encoded in such a way that it does not occupy any additional space. ABRIDGE exploits the sorted file order to store the difference between adjoining mapping positions further reducing space demand. It also discards read names for single-ended uniquely mapped reads which improves compression further. Finally, column-wise conversion of quality scores assists in achieving the best compression.

For ABRIDGE to be a viable compression software, the decompression should be achieved at an acceptable pace. ABRIDGE (with 7z compression) outperforms SAMCOMP, GENOZIP and DEEZ in terms of the duration for decompressing a lossless compressed SAM file. While ABRIDGE with ZPAQ attains the best compression it also takes a much higher time to decompress. Although the decompression time is less than downloading the fastq from NCBI and aligning it to the reference.

Our analysis establishes ABRIDGE as the most recent SAM alignment compressor that offers a very high compression ratio. For single-ended DNA-Seq reads, ABRIDGE produced a file which is 164 MB smaller than the next best compressor. This means that ABRIDGE can achieve an improvement of 15TB with 100K alignment files facilitating both storage and file transmission speed. Additionally, ABRIDGE compressed files can be randomly accessed making it convenient to perform searches without decompressing the entire file.

ABRIDGE provides users the option of choosing either lossless or lossy compression. It is recommended to use lossy compression in conjunction with downstream applications. For example, if the alignment files are produced for transcriptome assembly, then users can do away with quality scores and unmapped reads altogether. But if the downstream application involves SNP calling, then the quality scores (at least for the nucleotides that were a mismatch with the reference) should be preserved. Users are recommended to opt for ZPAQ if they choose to attain ultra high compression ratio. On the other hand, if decompression time is of essence then 7z compression would be the best choice. It is important to remember that ABRIDGE uses the reference file both for compression and decompression. Hence ABRIDGE stores a message digest of the reference to ensure that a correct copy is used for decompression.

An interesting future addition would be to expand ABRIDGE to other file types such as VCF, BED etc. Additionally, we would like to explore options to compress quality scores since those occupy the most space. Further, we will offer users the option to generate coverage information from compressed files directly. We are also collaborating with colleagues to adopt ABRIDGE as an acceptable file format to assembly and gene count software. We pledge to continually develop ABRIDGE to cater to a wide variety of file types storing biological information and facilitate its integration into existing pipelines.

## AVAILABILITY

ABRIDGE can be obtained from GitHub via this link https://github.com/sagnikbanerjee15/Abridge.

## ACKNOWLEDGEMENTS

The authors acknowledge Dr. Nathan Weeks for insightful discussion about release of software.

## SUPPLEMENTARY FIGURES

**Figure 3.**
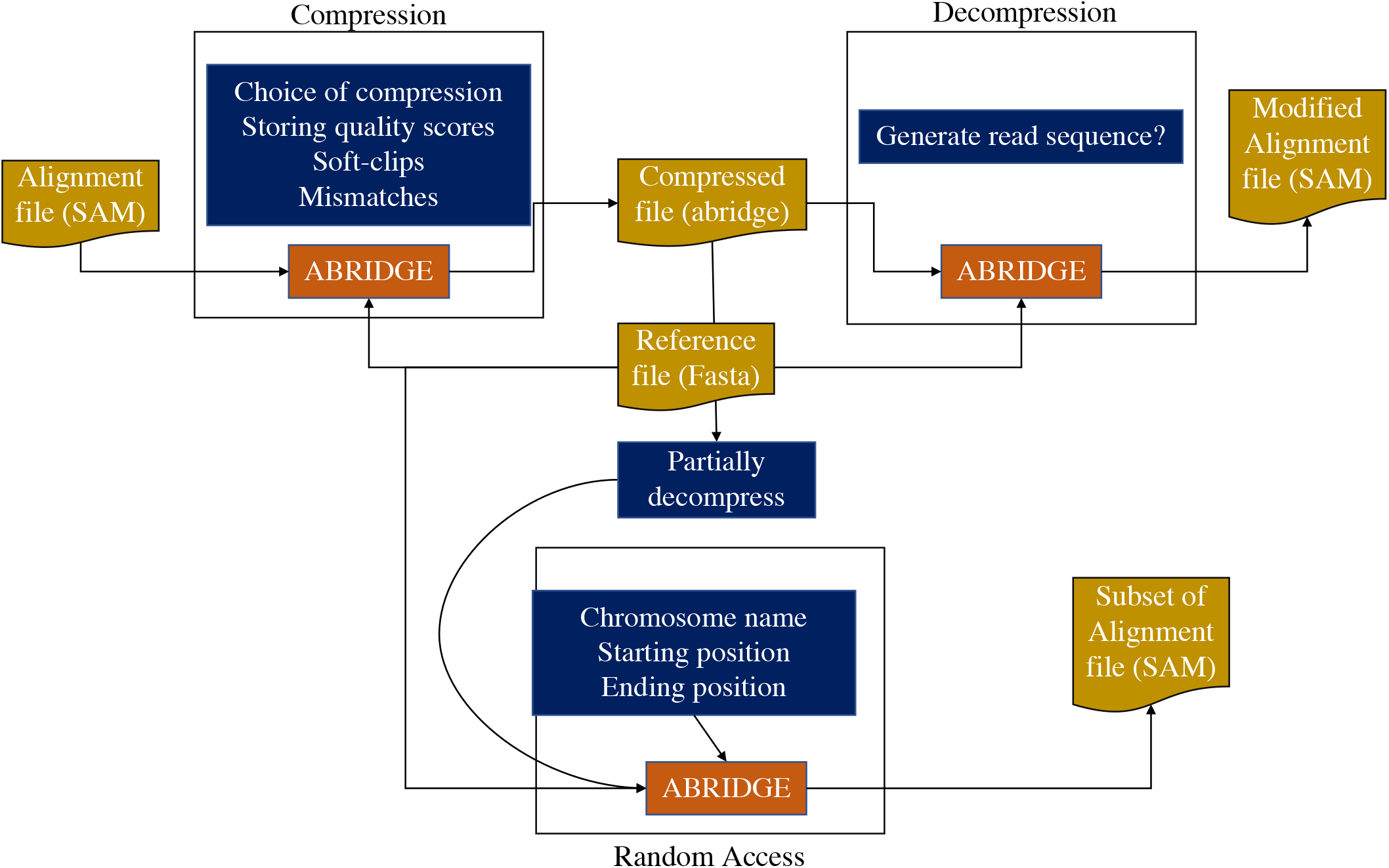
Overview of ABRIDGE software. ABRIDGE can be used for compressing alignment files in SAM format. Users have the option of providing multiple different modes of compression. The compressed file can be decompressed as and when required. ABRIDGE also offers users the opportunity to access random locations from the compressed file. All operations require the reference in fasta format.

**Figure 4.**
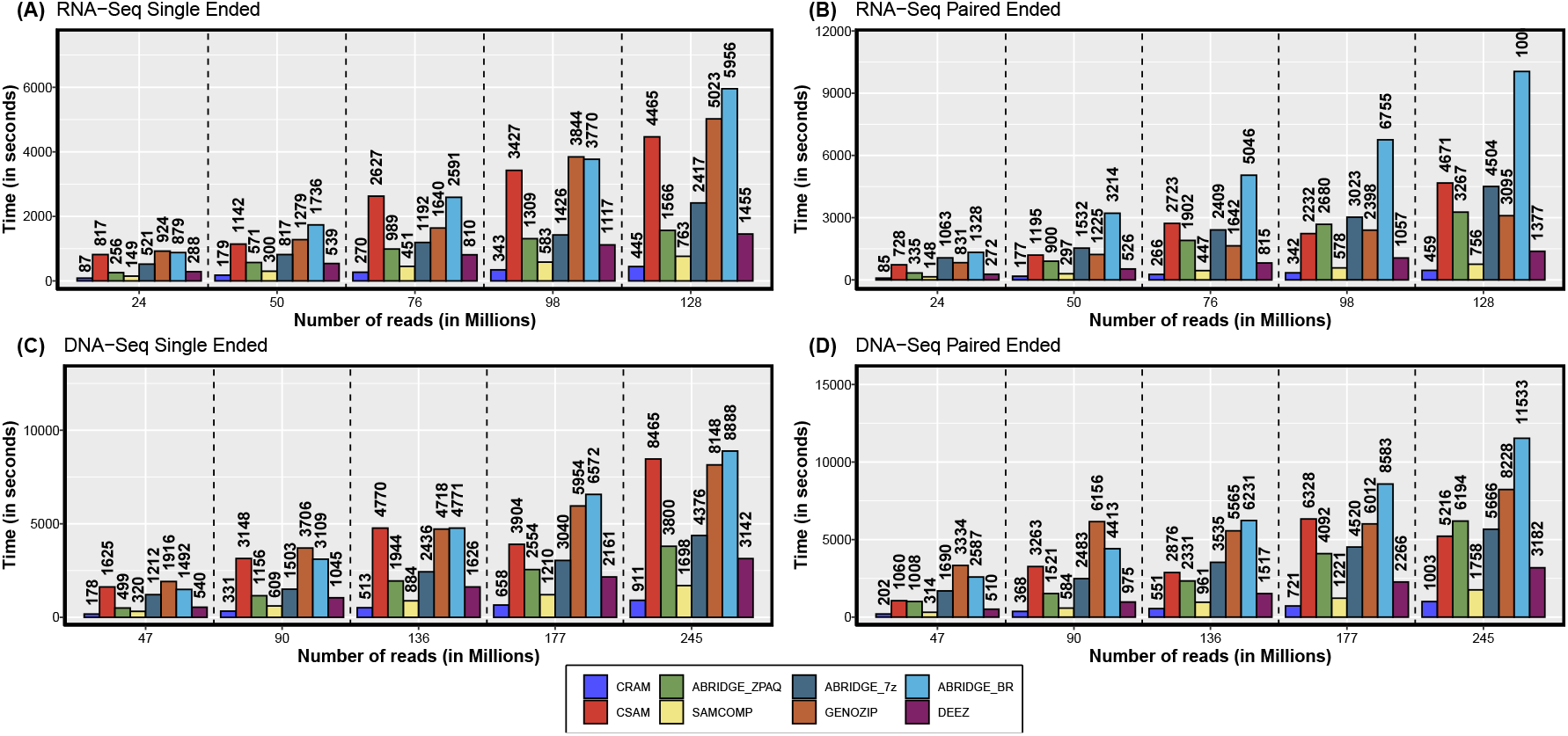
Comparison of time taken to compress SAM file.

**Figure 5.**
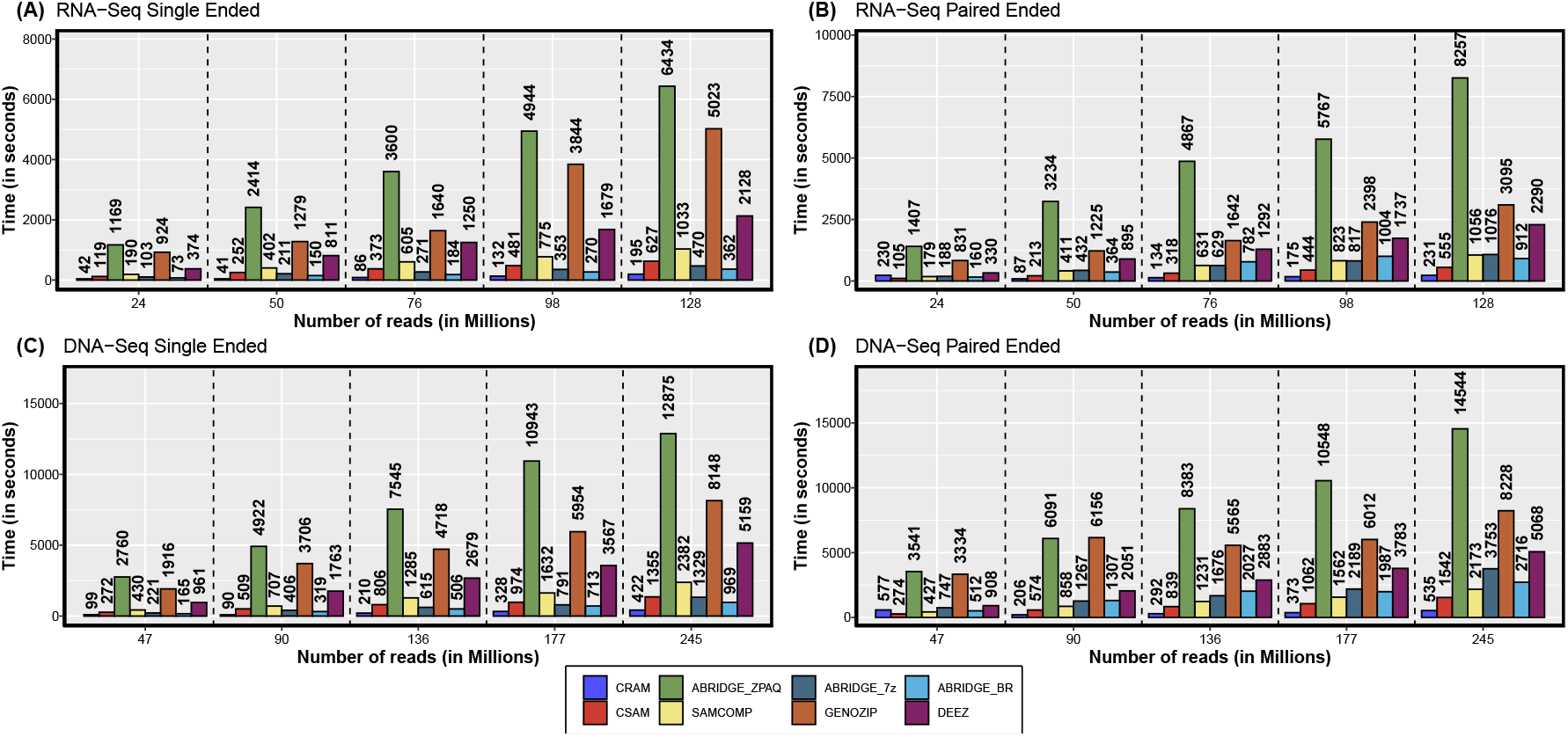
Comparison of time taken to decompress into SAM file.

**Figure 6.**
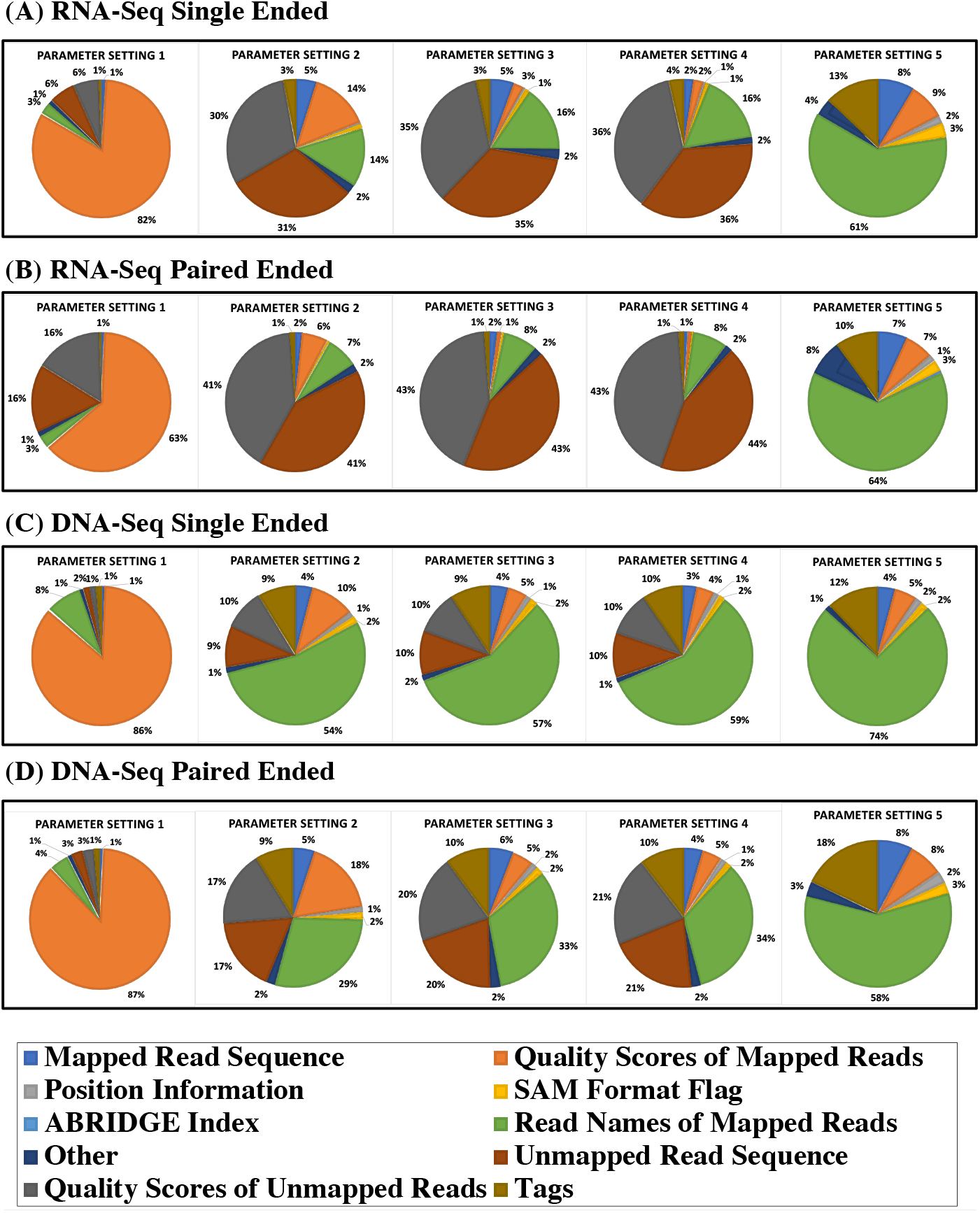
Various modes of compression offered by ABRIDGE.

**Figure 7.**
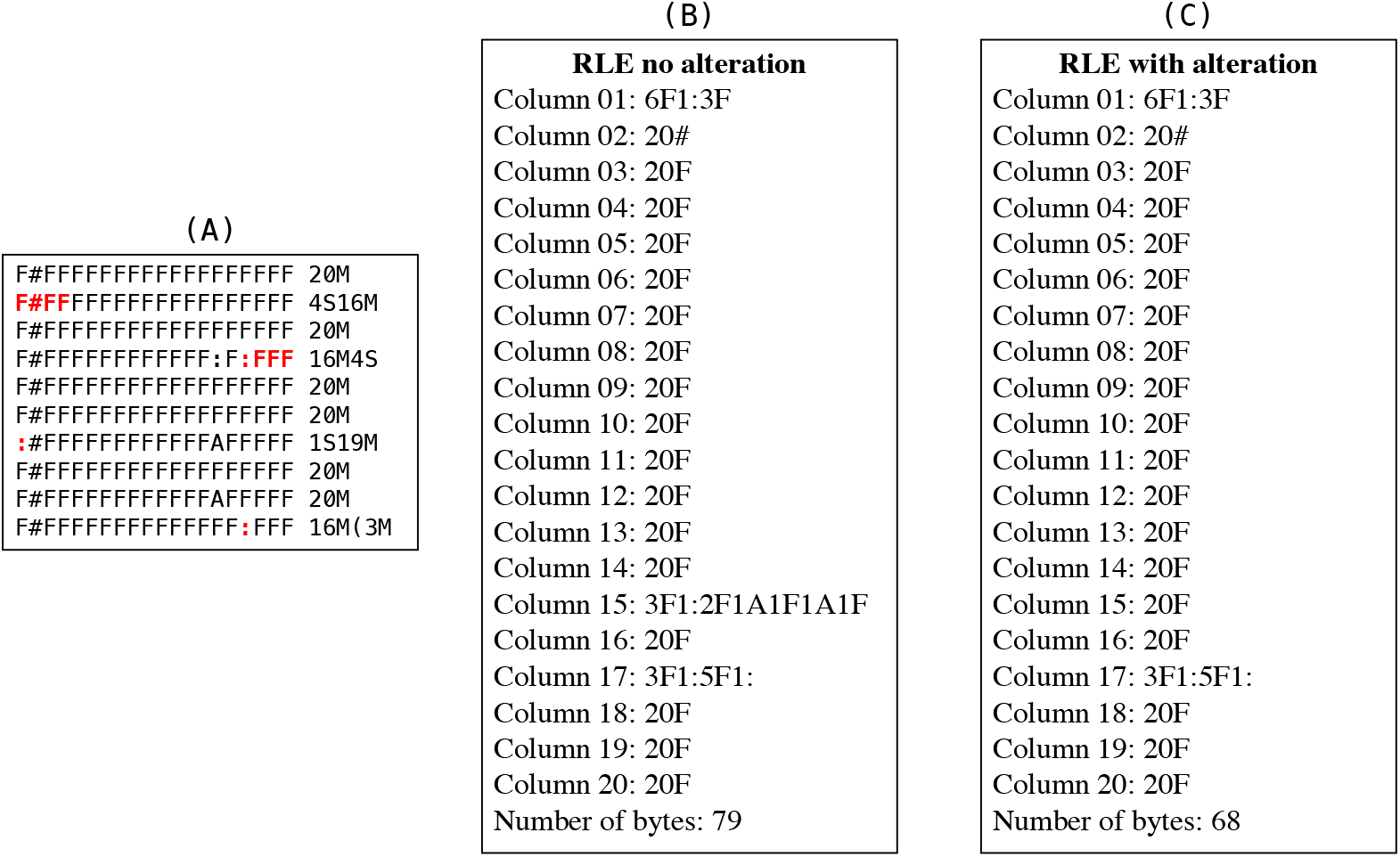
Run Length Encoding of columns in quality scores file. (A) Quality scores of first 20 bases of 10 alignments followed by the integrated CIGAR. Regions that are soft-clips or mismatches have been highlighted in Red. (B) Run length encoding of the quality scores without making any alteration. (C) Run length encoding by altering the quality scores of matched nucleotide bases

## SUPPLEMENTARY TABLES

**Table 2.**
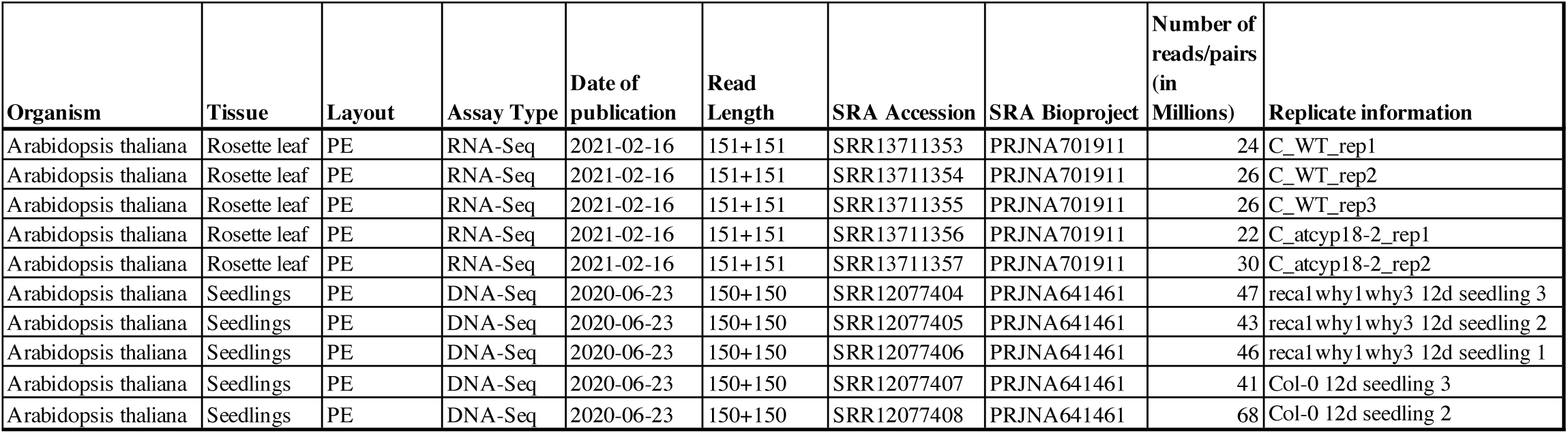
RNA-Seq and DNA-Seq samples for comparison.

**Table 3.**
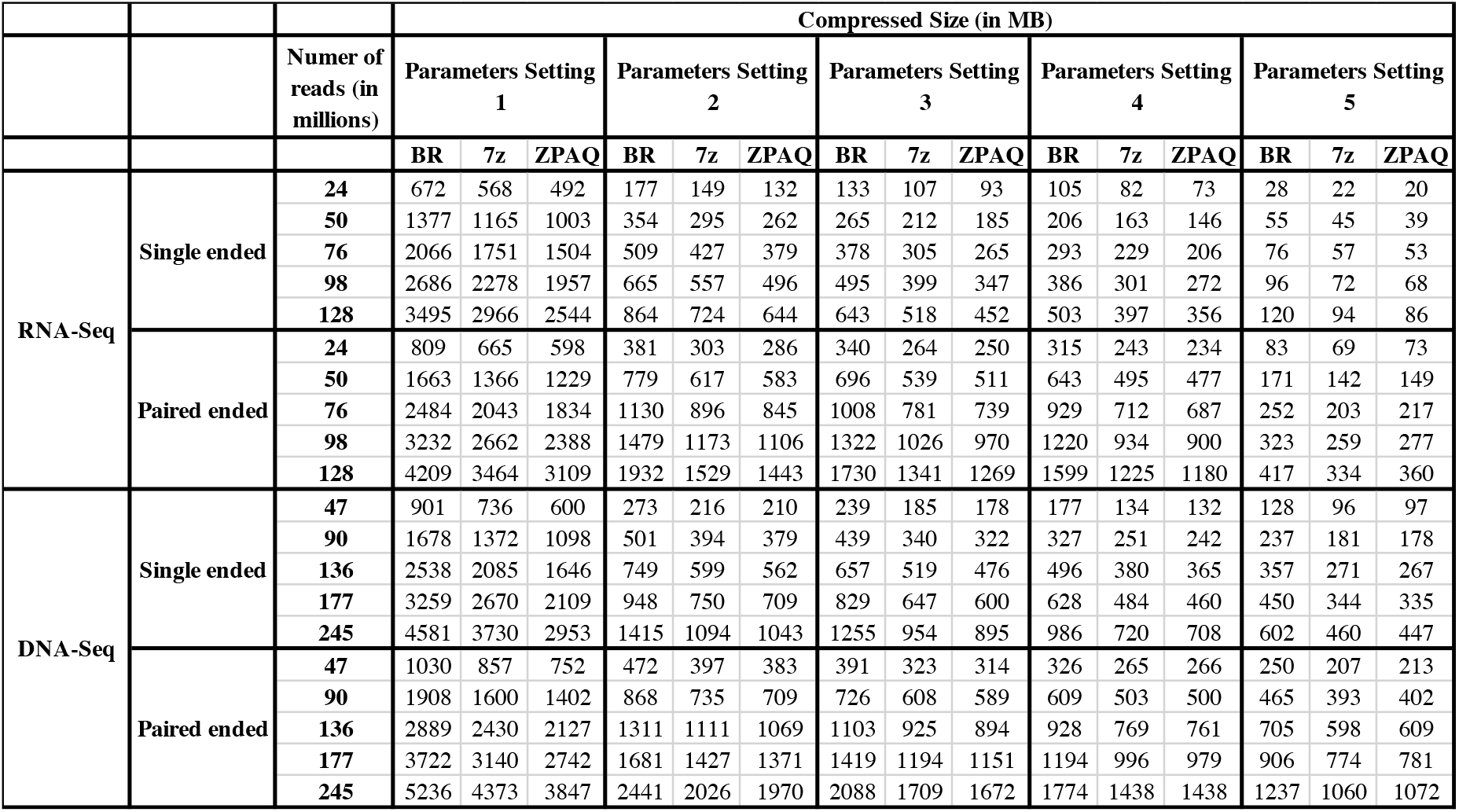
Compression achieved by ABRIDGE with different parameters and with different compressors.

**Table 4.**
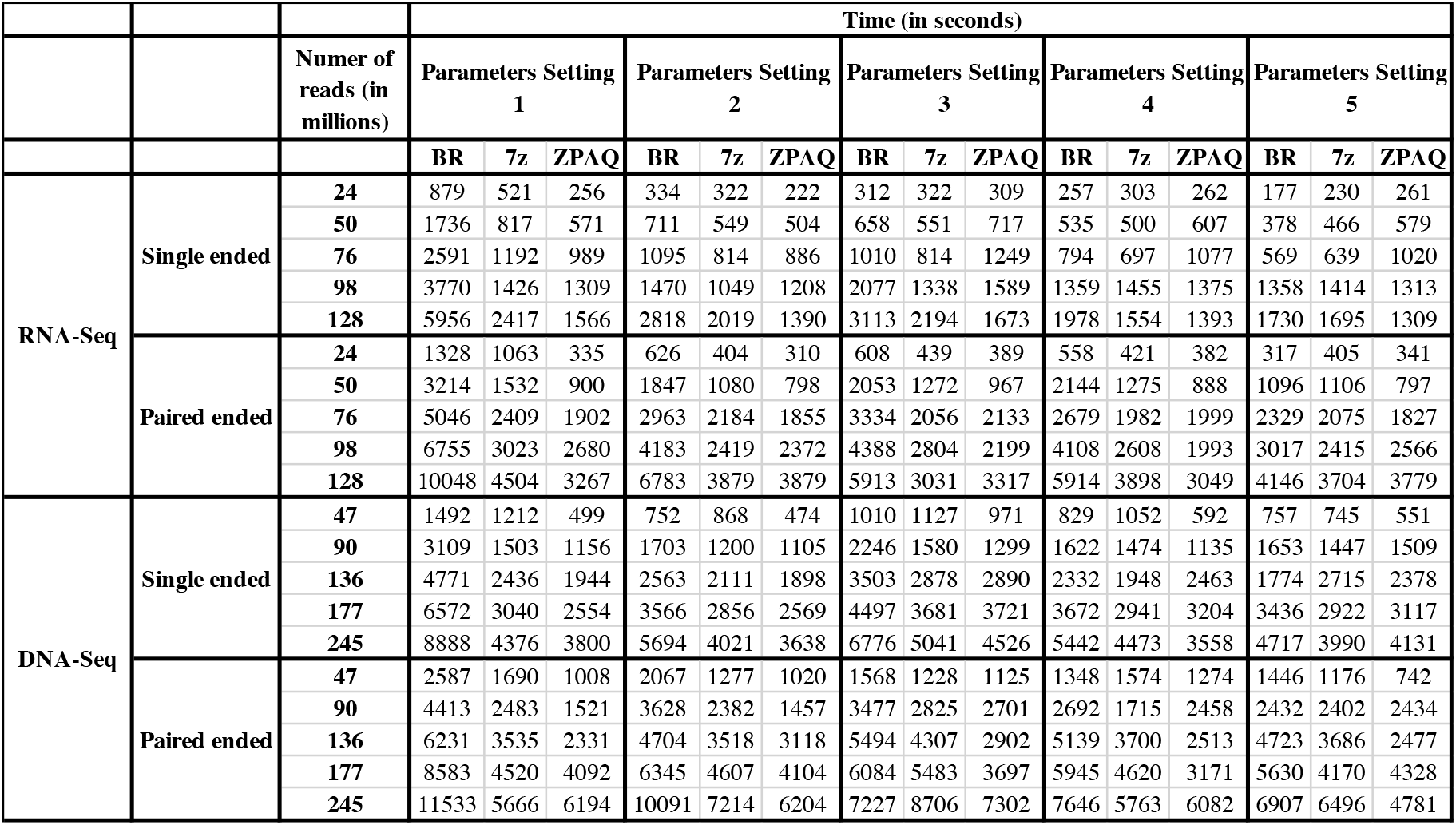
Duration of compression by ABRIDGE with different parameters and with different compressors.

**Table 5.**
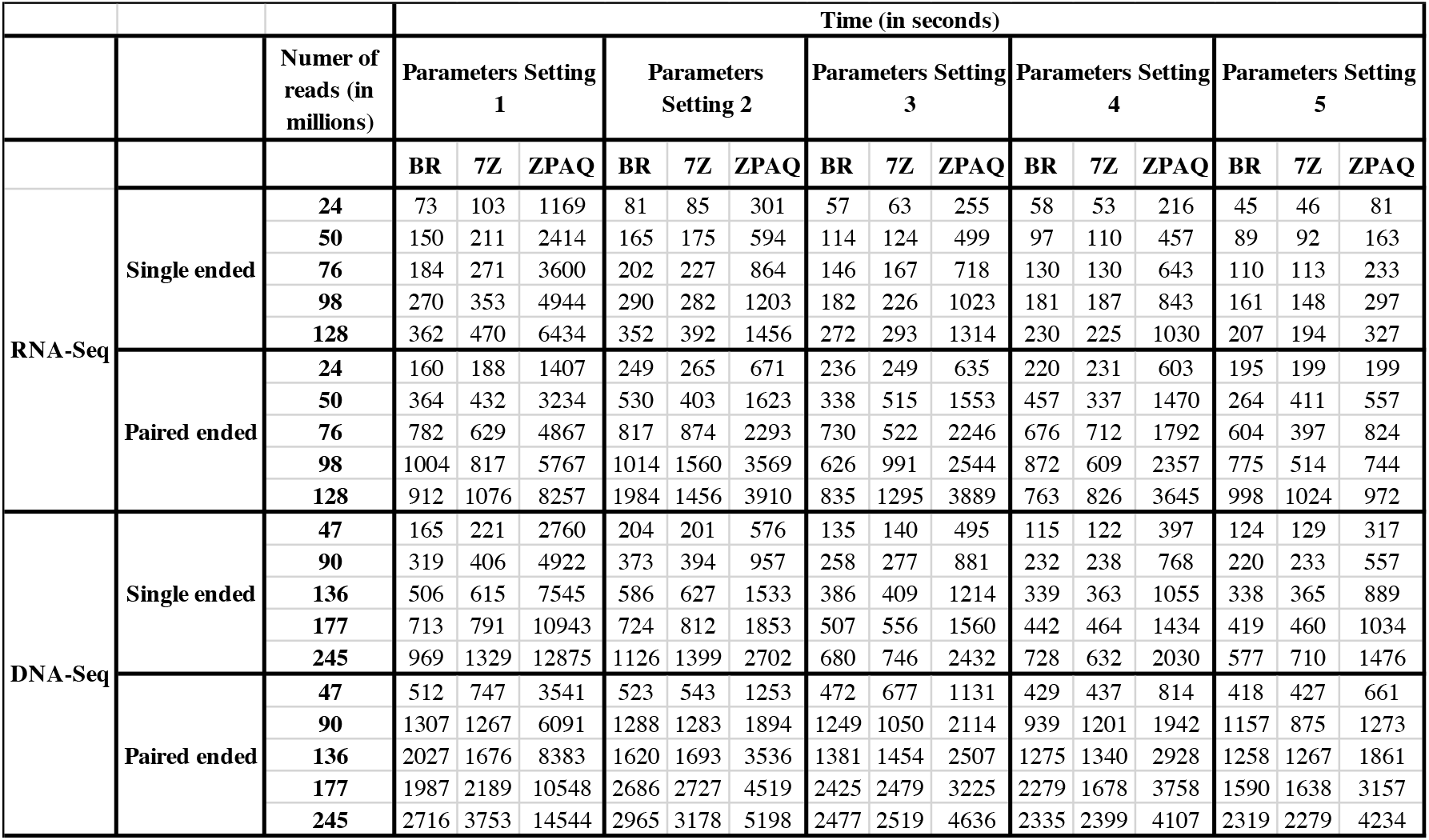
Duration of decompression by ABRIDGE with different parameters and with different compressors.

**Table 6.**
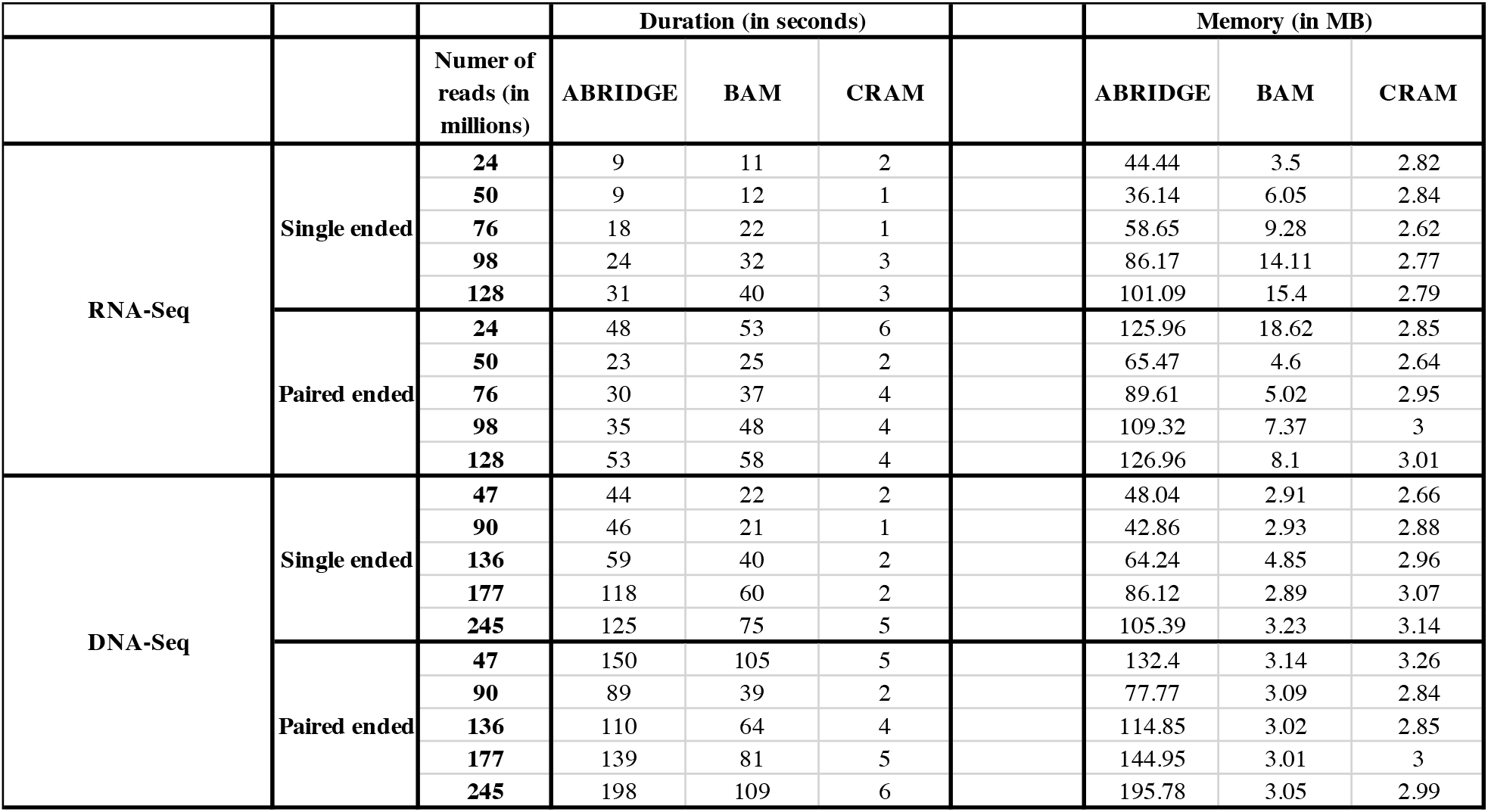
Duration and memory consumption of ABRIDGE to create index for random search.

**Table 7.**
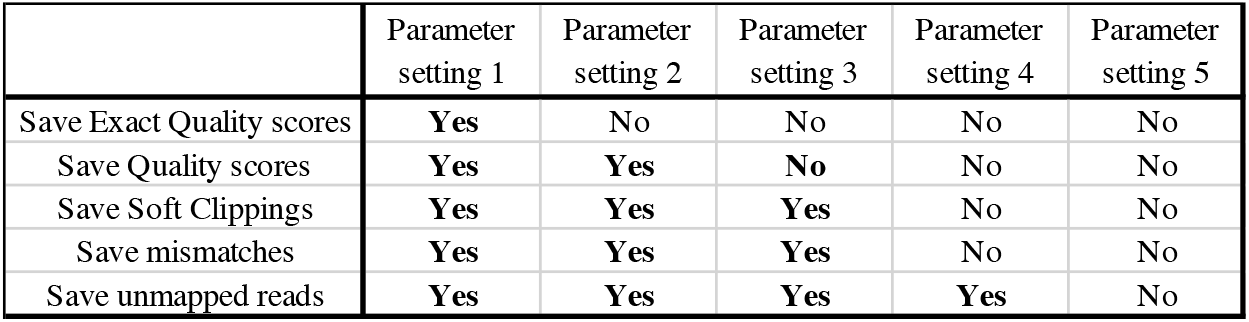
Illustration of arguments provided to ABRIDGE for each parameter setting.

**Table 8.**
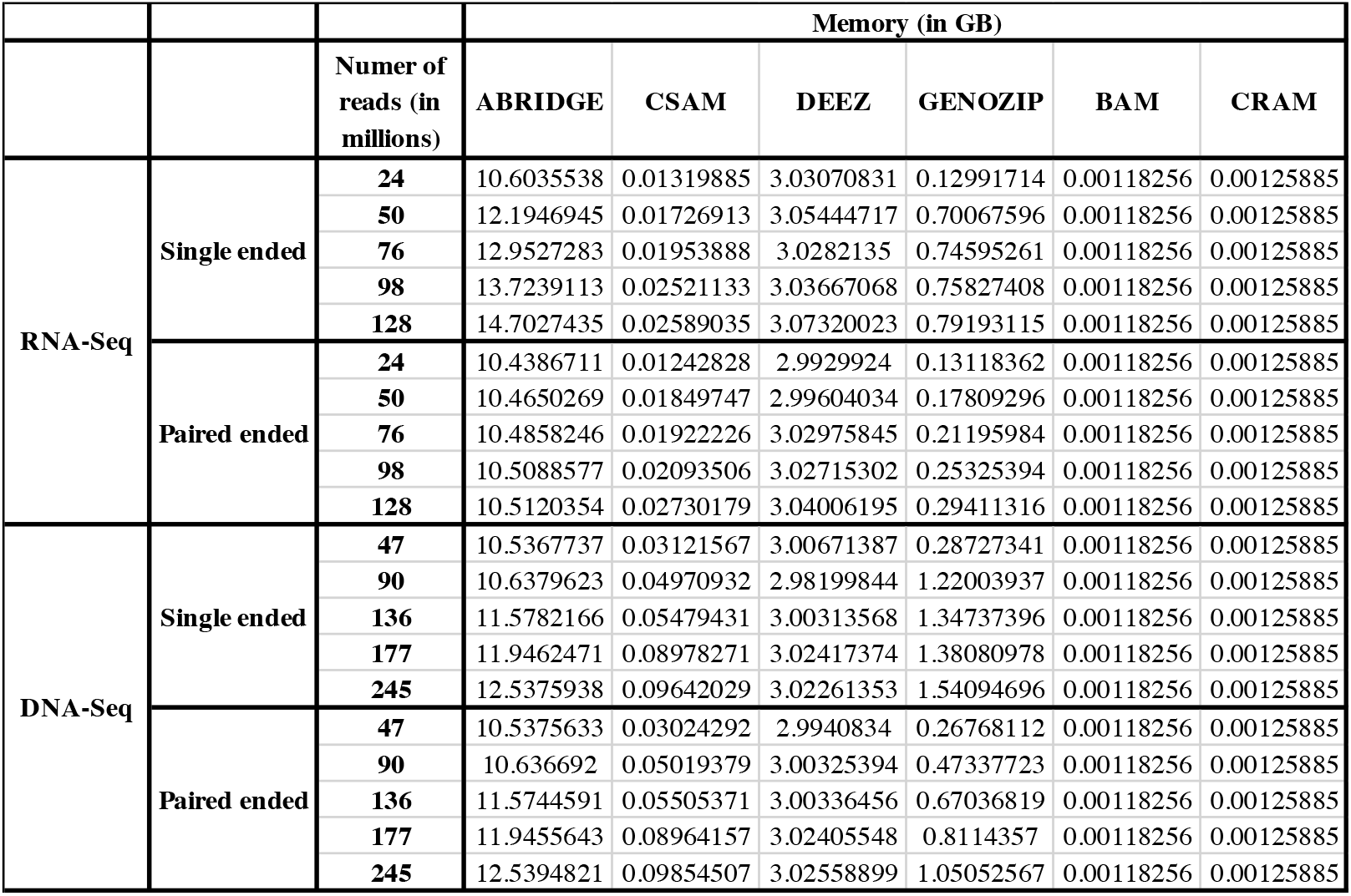
Memory consumed by different software while accessing a random location.

**Table 9.**
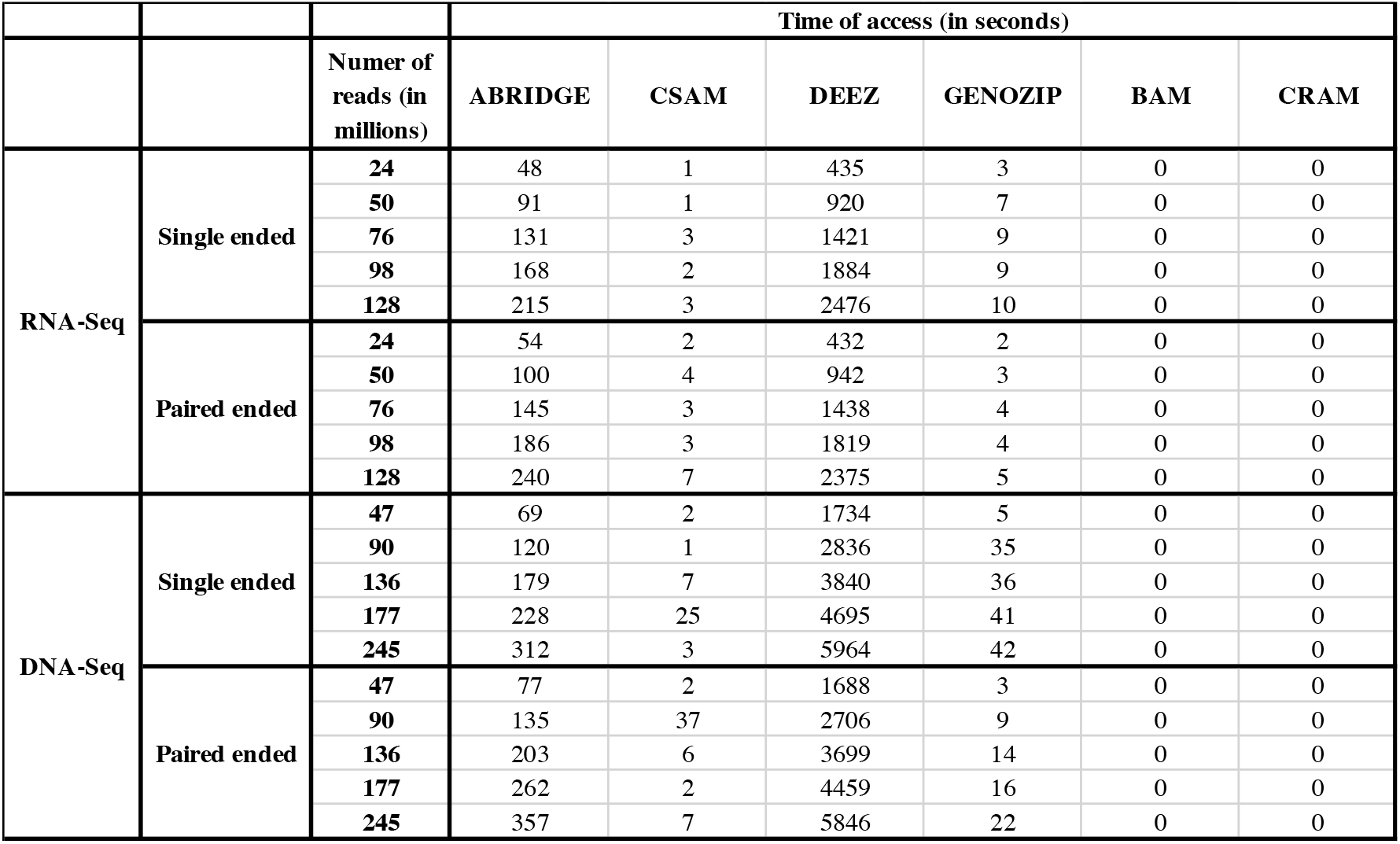
Duration of different software to access a random location.

## REFERENCES

1. Richard Hickman, Marcel C Van Verk, Anja J H Van Dijken, Marciel Pereira Mendes, Irene A Vroegop-Vos, Lotte Caarls, Merel Steenbergen, Ivo Van Der Nagel, Gert Jan Wesselink, and Aleksey Jironkin. Architecture and dynamics of the jasmonic acid gene regulatory network. The Plant Cell Online, pages tpc–00958, 2017.

2. Marie-Laure Erffelinck, Bianca Ribeiro, Maria Perassolo, Laurens Pauwels, Jacob Pollier, Veronique Storme, and Alain Goossens. A user-friendly platform for yeast two-hybrid library screening using next generation sequencing. PLOS ONE, 13(12):e0201270. 5 2018.

3. Matt Hunt, Sagnik Banerjee, Priyanka Surana, Meiling Liu, Greg Fuerst, Sandra Mathioni, Blake C Meyers, Dan Nettleton, and Roger P Wise. Small RNA discovery in the interaction between barley and the powdery mildew pathogen. BMC genomics, 20(1):610, 2019.

4. Manjula G. Elmore, Sagnik Banerjee, Kerry F. Pedley, Amy Ruck, and Steven A. Whitham. De novo transcriptome of Phakopsora pachyrhizi uncovers putative effector repertoire during infection. Physiological and Molecular Plant Pathology, 110, 4 2020.

5. Zhong Wang, Mark Gerstein, and Michael Snyder. RNA-Seq: a revolutionary tool for transcriptomics. Nature reviews genetics, 10(1):57–63, 2009.

6. Jason D Buenrostro, Beijing Wu, Howard Y Chang, and William J Greenleaf. ATAC-seq: a method for assaying chromatin accessibility genome-wide. Current protocols in molecular biology, 109(1):21–29, 2015.

7. Brian J Haas, Alexie Papanicolaou, Moran Yassour, Manfred Grabherr, Philip D Blood, Joshua Bowden, Matthew Brian Couger, David Eccles, Bo Li, Matthias Lieber, Matthew D MacManes, Michael Ott, Joshua Orvis, Nathalie Pochet, Francesco Strozzi, Nathan Weeks, Rick Westerman, Thomas William, Colin N Dewey, Robert Henschel, Richard D LeDuc, Nir Friedman, and Aviv Regev. De novo transcript sequence reconstruction from RNA-seq using the Trinity platform for reference generation and analysis. Nature protocols, 8(8):1494, 8 2013.

8. Mingfu Shao and Carl Kingsford. Accurate assembly of transcripts through phase-preserving graph decomposition. Nature Biotechnology, 35(12):1167–1169, 11 2017.

9. Sam Kovaka, Aleksey V Zimin, Geo M Pertea, Roham Razaghi, Steven L Salzberg, and Mihaela Pertea. Transcriptome assembly from long-read RNA-seq alignments with StringTie2. Genome Biology, 20(1):1–13, 2019.

10. Li Song, Sarven Sabunciyan, Guangyu Yang, and Liliana Florea. A multi-sample approach increases the accuracy of transcript assembly. Nature Communications, 10(1):5000, 12 2019.

11. Brian J Haas, Arthur L Delcher, Stephen M Mount, Jennifer R Wortman, Roger K Smith Jr, Linda I Hannick, Rama Maiti, Catherine M Ronning, Douglas B Rusch, and Christopher D Town. Improving the Arabidopsis genome annotation using maximal transcript alignment assemblies. Nucleic acids research, 31(19):5654–5666, 2003.

12. Carson Holt and Mark Yandell. MAKER2: an annotation pipeline and genome-database management tool for second-generation genome projects. BMC bioinformatics, 12(1):491, 12 2011.

13. Tomas Bruna, Katharina Hoff, Mario Stanke, Alexandre Lomsadze, and Mark Borodovsky. BRAKER2: Automatic Eukaryotic Genome Annotation with GeneMark-EP+ and AUGUSTUS Supported by a Protein Database. bioRxiv, 2020.

14. Sagnik Banerjee, Priyanka Bhandary, Margaret Woodhouse, Taner Z Sen, Roger P Wise, and Carson M Andorf. FINDER: An automated software package to annotate eukaryotic genes from RNA-Seq data and associated protein sequences. BMC bioinformatics, page 2021.02.04.429837, 4 2021.

15. Mark D Robinson, Davis J McCarthy, and Gordon K Smyth. edgeR: a Bioconductor package for differential expression analysis of digital gene expression data. Bioinformatics, 26(1):139–140, 2010.

16. Michael Love, Simon Anders, and Wolfgang Huber. Differential analysis of count data–the DESeq2 package. Genome Biology, 15:550, 2014.

17. Sagnik Banerjee, Subhadip Basu, and Mita Nasipuri. Big Data Analytics and Its Prospects in Computational Proteomics. In Information Systems Design and Intelligent Applications, volume 340, pages 591–598. Springer, 2015.

18. Sagnik Banerjee, Valeria Velasquez-Zapata, Gregory Fuerst, J Mitch Elmore, and Roger P Wise. NGPINT: A Next-generation protein-protein interaction software. Breifings in Bioinformatics, 22:in press, 1 2021.

19. Valeria Velásquez-Zapata, James Mitch Elmore, Sagnik Banerjee, Karin S Dorman, and Roger P Wise. Y2H-SCORES: A statistical framework to infer protein-protein interactions from next-generation yeast-two-hybrid sequence data. bioarxiv, 2020.

20. Heng Li, Bob Handsaker, Alec Wysoker, Tim Fennell, Jue Ruan, Nils Homer, Gabor Marth, Goncalo Abecasis, and Richard Durbin. The sequence alignment/map format and SAMtools. Bioinformatics, 25(16):2078–2079, 2009.

21. Markus Hsi-Yang Fritz, Rasko Leinonen, Guy Cochrane, and Ewan Birney. Efficient storage of high throughput DNA sequencing data using reference-based compression. Genome research, 21(5):734–740, 2011.

22. Alexander Dobin, Carrie A Davis, Felix Schlesinger, Jorg Drenkow, Chris Zaleski, Sonali Jha, Philippe Batut, Mark Chaisson, and Thomas R Gingeras. STAR: Ultrafast universal RNA-seq aligner. Bioinformatics, 29(1):15–21, 2013.

23. Alexander Dobin, Thomas R Gingeras, Cold Spring, Roberto Flores, Joshua Sampson, Rob Knight, Nicholas Chia, and High-throughput Sequencing Technologies. Mapping RNA-seq with STAR. Curr Protoc Bioinformatics, 51(4):586–597, 2016.

24. Daehwan Kim, Ben Langmead, and Steven L Salzberg. HISAT: a fast spliced aligner with low memory requirements. Nature Methods, 12(4):357–360, 3 2015.

25. José M Abúin, Juan C Pichel, Tomás F Pena, and Jorge Amigo. BigBWA: approaching the Burrows–Wheeler aligner to Big Data technologies. Bioinformatics, page btv506, 2015.

26. Raffaele Giancarlo, Simona E Rombo, and Filippo Utro. Compressive biological sequence analysis and archival in the era of high-throughput sequencing technologies. Briefings in bioinformatics, 15(3):390–406, 2014.

27. Morteza Hosseini, Diogo Pratas, and Armando J Pinho. A survey on data compression methods for biological sequences. Information, 7(4):56, 2016.

28. Ibrahim Numanagić, James K Bonfield, Faraz Hach, Jan Voges, Jörn Ostermann, Claudio Alberti, Marco Mattavelli, and S Cenk Sahinalp. Comparison of high-throughput sequencing data compression tools. nature methods, 13(12):1005–1008, 2016.

29. Niko Popitsch and Arndt von Haeseler. NGC: lossless and lossy compression of aligned high-throughput sequencing data. Nucleic acids research, 41(1):e27–e27, 2013.

30. Faraz Hach, Ibrahim Numanagic, and S Cenk Sahinalp. DeeZ: reference-based compression by local assembly. Nature methods, 11(11):1082– 1084, 2014.

31. Divon Lan, Ray Tobler, Yassine Souilmi, and Bastien Llamas. Genozip-A Universal Extensible Genomic Data Compressor. Bioinformatics, 2021.

32. Daniel C Jones, Walter L Ruzzo, Xinxia Peng, and Michael G Katze. Compression of next-generation sequencing reads aided by highly efficient de novo assembly. Nucleic acids research, 40(22):e171–e171, 2012.

33. Rodrigo Canóvas, Alistair Moffat, and Andrew Turpin. Csam: Compressed sam format. Bioinformatics, 32(24):3709–3716, 2016.

34. James K Bonfield and Matthew V Mahoney. Compression of FASTQ and SAM format sequencing data. PloS one, 8(3):e59190, 2013.

