## Supplementary document for "ABRIDGE: An ultra-compression software for SAM alignment files"

### SUPPLEMENTAL MATERIALS

#### SAM file format

We illustrate the various fields of SAM format below. The first 11 fields of the SAM format is mandatory while the rest are optional.

*Required fields* 1. **QNAME** (Type: String) - Name of the query template
2. **FLAG** (Type: Integer) - A format flag to denote the type of alignment
3. **RNAME** (Type: String) - The name of the reference
4. **POS** (Type: Integer) - The position where the short read is mapped
5. **MAPQ** (Type: Integer) - A score to denote the quality of alignment
6. **CIGAR** (Type: String) - Stands for Compressed Idiosyncratic Gapped Alignment Report. A string that outlines the matches, inserts and deletions is the alignment
7. **RNEXT** (Type: String) - Reference name to which the read in the other pair has mapped
8. **PNEXT** (Type: Integer) - Position to which the read in the other pair has mapped
9. **TLEN** (Type: Integer) - Length of the template
10. **SEQ** (Type: String) - The nucleotide sequence of the read
11. **QUAL** (Type: String) - The quality scores as reported by the sequencing machine

*Optional fields* A number of optional fields can be included in a SAM file. Among them, the most important are NH, MD and XS. All three tags are required by ABRIDGE.

#### Selection of samples for testing

A total of five RNA-Seq and five DNA-Seq samples were chosen from NCBI SRA for testing and for comparing all the compression software. All the samples chosen were paired-ended and sequenced to 150 bp. Single-ended samples were generated by merging the two mate pairs together. We chose samples from different sequencing assays to demonstrate the superiority of ABRIDGE over other compression software across the entire spectrum. To demonstrate the linear increase of compressed size with increase in file size, samples were merged together to mimic deeply sequenced samples **Supplementary Table** .

#### Alignment to reference

We used STAR ((**?**)) to align the short reads with a minimum mapping threshold of 75%. Even-though STAR is designed to align RNA-Seq reads we modified the settings to enforce STAR to map DNA-Seq reads without any splices. For DNA-Seq reads mapping was carried out with the following parameters - `--scoreGap -100, --scoreGapNoncan -100, --scoreGapGCAG -100, --scoreGapATAC -100 and --alignSJoverhangMin 500`. The intentional high penalty for splice generation forced all the alignments to be unspliced. All the alignments of DNA-Seq samples were inspected to ensure that there were no spliced alignments present.

#### ABRIDGE Compression

ABRIDGE performs compression in two steps. During the first step, ABRIDGE restructures the data to discard any redundant information. Since the alignment file is always accepted in sorted format, ABRIDGE stores only the difference between consecutive mapped positions. For a deeply sequenced sample, this helps save a lot of space. Nucleotide sequences are entirely eliminated except for mismatches and indels. The sequences can be later reconstructed from the reference and the position stored in the compressed file. Unlike nucleotide sequence, quality scores cannot be ”mapped” to any reference. Hence, all quality scores need to be stored if and when the user requests for it. This can lead to a rise in the space demand. Hence, we employed a different method to compress quality scores.

*Run Length encoding of quality scores* To compress quality scores, we implement run length encoding. Instead of encoding quality scores of each read, we apply run-length encoding simultaneously for all quality scores in a particular nucleotide position of the read. If the user allows ABRIDGE to modify the quality scores of matched bases, then the software achieves even better compression.

#### Commands to run other software

In this section we discuss the commands used to generate compressed files by DEEZ, SAMCOMP, GENOZIP and CSAM. DEEZ was executed with lossy values of `0, 50 and 99` and both modes to encode quality. SAMCOMP was executed only with the sorted alignment file. Compression using GENOZIP was performed by both modes of compression and was launched with the ‘optimize‘ parameter. CSAM was also launched in both lossy and lossless modes.

#### Future Work

ABRIDGE will be updated in the future to compress other types of files that store biological information like BED, VCF, etc. To enhance compression, we will further explore other techniques to compress quality scores. Currently, ABRIDGE stores the read names for paired ended reads and also for multi-mapped single ended reads. We will modify our algorithm to retain all relevant information without having to store read names.

### SUPPLEMENTARY FIGURES

### SUPPLEMENTARY TABLES

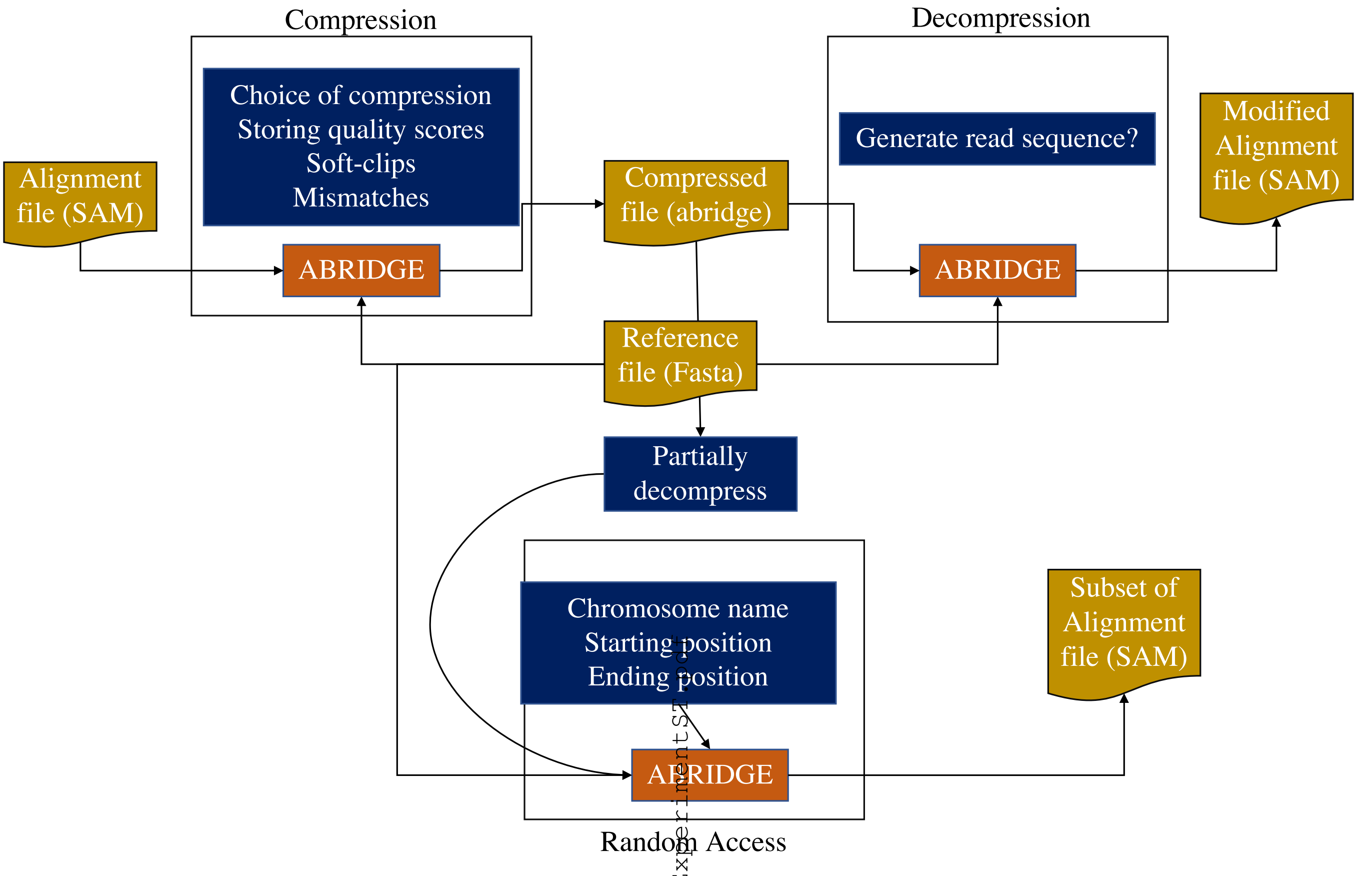


**Figure 1. Overview of ABRIDGE software** ABRIDGE can be used for compressing alignment files in SAM format. Users have the option of providing multiple different modes of compression. The compressed file can be decompressed as and when required. ABRIDGE also offers users the opportunity to access random locations from the compressed file. All operations require the reference in fasta format.

images/abridge_ListOfNCBI-SRASamplesForExperimentSI.pdf

**Table 1. RNA-Seq and DNA-Seq samples for comparison**

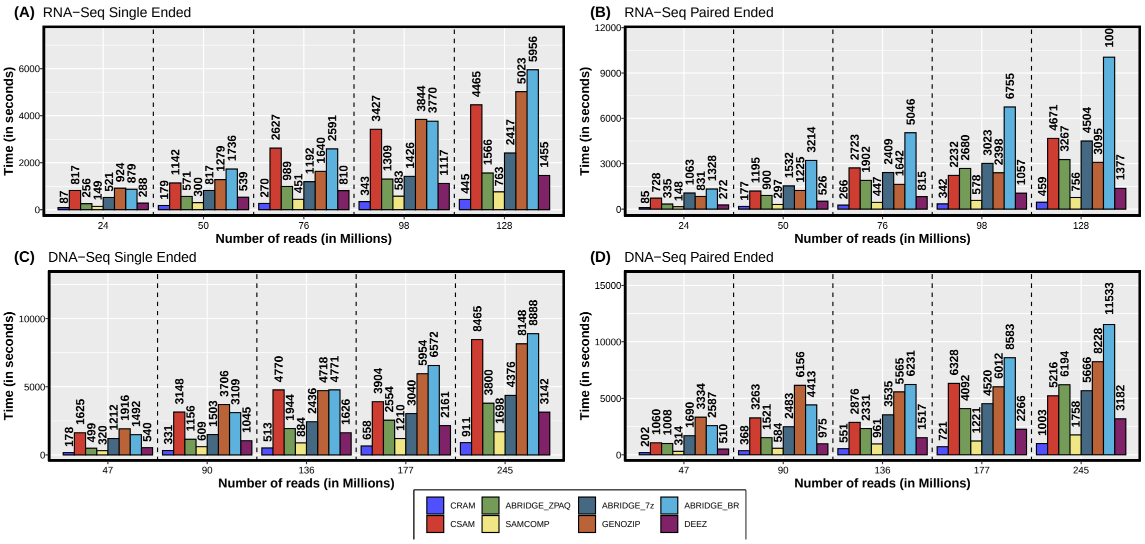


**Figure 2. Comparison of time taken to compress SAM file**

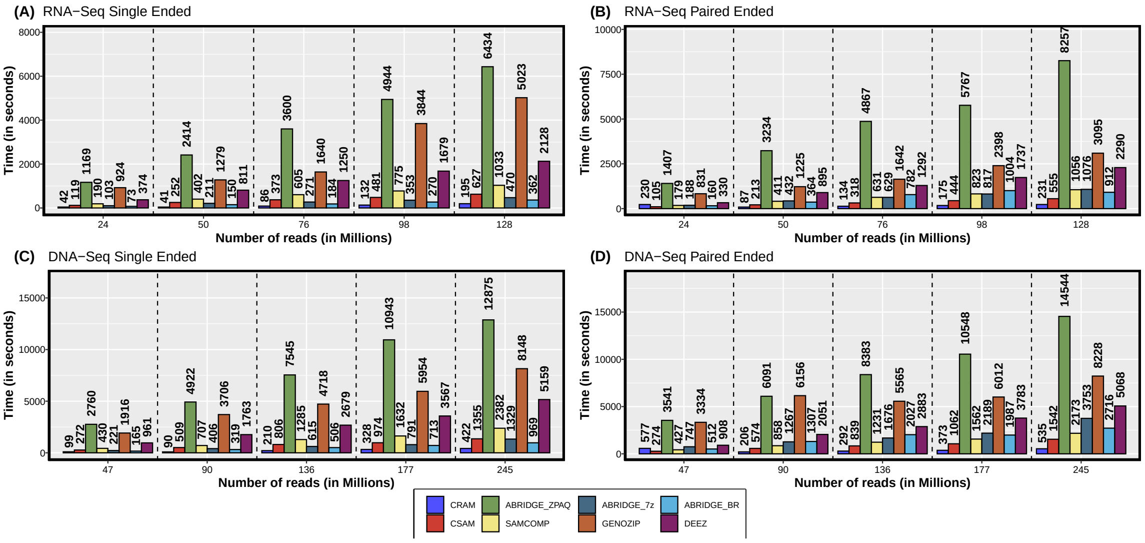


**Figure 3. Comparison of time taken to decompress into SAM file**

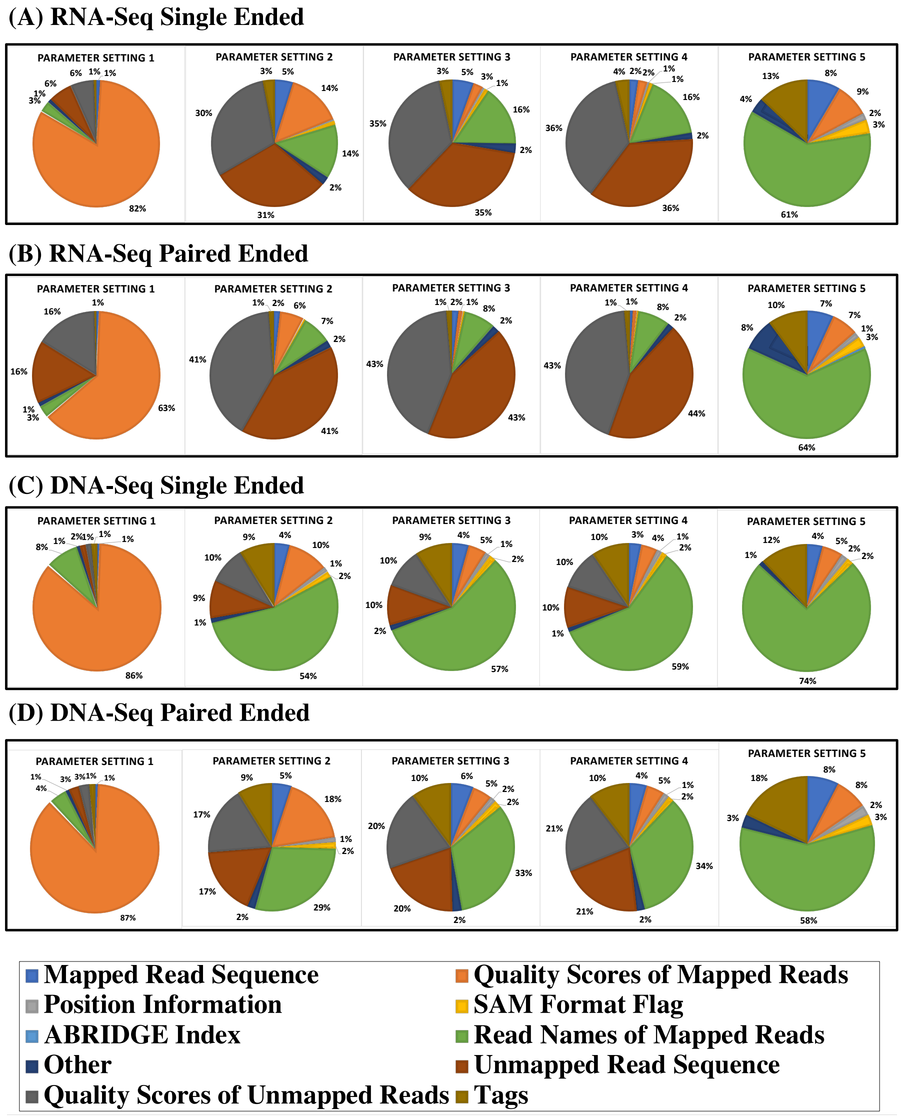


**Figure 4.** **Various modes of compression offered by ABRIDGE**

(A)

```
F#FFFFFFFFFFFFFFFFFF 20M
F#FFFFFFFFFFFFFFFFFF 4S16M
F#FFFFFFFFFFFFFFFFFF 20M
F#FFFFFFFFFFFF:F:FFF 16M4S
F#FFFFFFFFFFFFFFFFFF 20M
F#FFFFFFFFFFFFFFFFFF 20M
:#FFFFFFFFFFFFAFFFFF 1S19M
F#FFFFFFFFFFFFFFFFFF 20M
F#FFFFFFFFFFFFAFFFFF 20M
F#FFFFFFFFFFFFFF:FFF 16M(3M
```

(B)

**RLE no alteration**
Column 01: 6F1:3F
Column 02: 20#
Column 03: 20F
Column 04: 20F
Column 05: 20F
Column 06: 20F
Column 07: 20F
Column 08: 20F
Column 09: 20F
Column 10: 20F
Column 11: 20F
Column 12: 20F
Column 13: 20F
Column 14: 20F
Column 15: 3F1:2F1A1F1A1F
Column 16: 20F
Column 17: 3F1:5F1:
Column 18: 20F
Column 19: 20F
Column 20: 20F
Number of bytes: 79

(C)

**RLE with alteration**
Column 01: 6F1:3F
Column 02: 20#
Column 03: 20F
Column 04: 20F
Column 05: 20F
Column 06: 20F
Column 07: 20F
Column 08: 20F
Column 09: 20F
Column 10: 20F
Column 11: 20F
Column 12: 20F
Column 13: 20F
Column 14: 20F
Column 15: 20F
Column 16: 20F
Column 17: 3F1:5F1:
Column 18: 20F
Column 19: 20F
Column 20: 20F
Number of bytes: 68

**Figure 5. Run Length Encoding of columns in quality scores file** (A) Quality scores of first 20 bases of 10 alignments followed by the integrated CIGAR. Regions that are soft-clips or mismatches have been highlighted in Red. (B) Run length encoding of the quality scores without making any alteration. (C) Run length encoding by altering the quality scores of matched nucleotide bases

| | | | **Compressed Size (in MB)** | | | | | | | | | | | | | | |
|---|---|---|---|---|---|---|---|---|---|---|---|---|---|---|---|---|---|
| | | **Numer of reads (in millions)** | **Parameters Setting 1** | | | **Parameters Setting 2** | | | **Parameters Setting 3** | | | **Parameters Setting 4** | | | **Parameters Setting 5** | | |
| | | | **BR** | **7z** | **ZPAQ** | **BR** | **7z** | **ZPAQ** | **BR** | **7z** | **ZPAQ** | **BR** | **7z** | **ZPAQ** | **BR** | **7z** | **ZPAQ** |
| **RNA-Seq** | **Single ended** | **24** | 672 | 568 | 492 | 177 | 149 | 132 | 133 | 107 | 93 | 105 | 82 | 73 | 28 | 22 | 20 |
| | | **50** | 1377 | 1165 | 1003 | 354 | 295 | 262 | 265 | 212 | 185 | 206 | 163 | 146 | 55 | 45 | 39 |
| | | **76** | 2066 | 1751 | 1504 | 509 | 427 | 379 | 378 | 305 | 265 | 293 | 229 | 206 | 76 | 57 | 53 |
| | | **98** | 2686 | 2278 | 1957 | 665 | 557 | 496 | 495 | 399 | 347 | 386 | 301 | 272 | 96 | 72 | 68 |
| | | **128** | 3495 | 2966 | 2544 | 864 | 724 | 644 | 643 | 518 | 452 | 503 | 397 | 356 | 120 | 94 | 86 |
| | **Paired ended** | **24** | 809 | 665 | 598 | 381 | 303 | 286 | 340 | 264 | 250 | 315 | 243 | 234 | 83 | 69 | 73 |
| | | **50** | 1663 | 1366 | 1229 | 779 | 617 | 583 | 696 | 539 | 511 | 643 | 495 | 477 | 171 | 142 | 149 |
| | | **76** | 2484 | 2043 | 1834 | 1130 | 896 | 845 | 1008 | 781 | 739 | 929 | 712 | 687 | 252 | 203 | 217 |
| | | **98** | 3232 | 2662 | 2388 | 1479 | 1173 | 1106 | 1322 | 1026 | 970 | 1220 | 934 | 900 | 323 | 259 | 277 |
| | | **128** | 4209 | 3464 | 3109 | 1932 | 1529 | 1443 | 1730 | 1341 | 1269 | 1599 | 1225 | 1180 | 417 | 334 | 360 |
| **DNA-Seq** | **Single ended** | **47** | 901 | 736 | 600 | 273 | 216 | 210 | 239 | 185 | 178 | 177 | 134 | 132 | 128 | 96 | 97 |
| | | **90** | 1678 | 1372 | 1098 | 501 | 394 | 379 | 439 | 340 | 322 | 327 | 251 | 242 | 237 | 181 | 178 |
| | | **136** | 2538 | 2085 | 1646 | 749 | 599 | 562 | 657 | 519 | 476 | 496 | 380 | 365 | 357 | 271 | 267 |
| | | **177** | 3259 | 2670 | 2109 | 948 | 750 | 709 | 829 | 647 | 600 | 628 | 484 | 460 | 450 | 344 | 335 |
| | | **245** | 4581 | 3730 | 2953 | 1415 | 1094 | 1043 | 1255 | 954 | 895 | 986 | 720 | 708 | 602 | 460 | 447 |
| | **Paired ended** | **47** | 1030 | 857 | 752 | 472 | 397 | 383 | 391 | 323 | 314 | 326 | 265 | 266 | 250 | 207 | 213 |
| | | **90** | 1908 | 1600 | 1402 | 868 | 735 | 709 | 726 | 608 | 589 | 609 | 503 | 500 | 465 | 393 | 402 |
| | | **136** | 2889 | 2430 | 2127 | 1311 | 1111 | 1069 | 1103 | 925 | 894 | 928 | 769 | 761 | 705 | 598 | 609 |
| | | **177** | 3722 | 3140 | 2742 | 1681 | 1427 | 1371 | 1419 | 1194 | 1151 | 1194 | 996 | 979 | 906 | 774 | 781 |
| | | **245** | 5236 | 4373 | 3847 | 2441 | 2026 | 1970 | 2088 | 1709 | 1672 | 1774 | 1438 | 1438 | 1237 | 1060 | 1072 |

**Table 2. Compression achieved by ABRIDGE with different parameters and with different compressors**

| | | | Time (in seconds) | | | | | | | | | | | | | | |
|---|---|---|---|---|---|---|---|---|---|---|---|---|---|---|---|---|---|
| | | **Numer of reads (in millions)** | **Parameters Setting 1** | | | **Parameters Setting 2** | | | **Parameters Setting 3** | | | **Parameters Setting 4** | | | **Parameters Setting 5** | | |
| | | | **BR** | **7z** | **ZPAQ** | **BR** | **7z** | **ZPAQ** | **BR** | **7z** | **ZPAQ** | **BR** | **7z** | **ZPAQ** | **BR** | **7z** | **ZPAQ** |
| **RNA-Seq** | **Single ended** | **24** | 879 | 521 | 256 | 334 | 322 | 222 | 312 | 322 | 309 | 257 | 303 | 262 | 177 | 230 | 261 |
| | | **50** | 1736 | 817 | 571 | 711 | 549 | 504 | 658 | 551 | 717 | 535 | 500 | 607 | 378 | 466 | 579 |
| | | **76** | 2591 | 1192 | 989 | 1095 | 814 | 886 | 1010 | 814 | 1249 | 794 | 697 | 1077 | 569 | 639 | 1020 |
| | | **98** | 3770 | 1426 | 1309 | 1470 | 1049 | 1208 | 2077 | 1338 | 1589 | 1359 | 1455 | 1375 | 1358 | 1414 | 1313 |
| | | **128** | 5956 | 2417 | 1566 | 2818 | 2019 | 1390 | 3113 | 2194 | 1673 | 1978 | 1554 | 1393 | 1730 | 1695 | 1309 |
| | **Paired ended** | **24** | 1328 | 1063 | 335 | 626 | 404 | 310 | 608 | 439 | 389 | 558 | 421 | 382 | 317 | 405 | 341 |
| | | **50** | 3214 | 1532 | 900 | 1847 | 1080 | 798 | 2053 | 1272 | 967 | 2144 | 1275 | 888 | 1096 | 1106 | 797 |
| | | **76** | 5046 | 2409 | 1902 | 2963 | 2184 | 1855 | 3334 | 2056 | 2133 | 2679 | 1982 | 1999 | 2329 | 2075 | 1827 |
| | | **98** | 6755 | 3023 | 2680 | 4183 | 2419 | 2372 | 4388 | 2804 | 2199 | 4108 | 2608 | 1993 | 3017 | 2415 | 2566 |
| | | **128** | 10048 | 4504 | 3267 | 6783 | 3879 | 3879 | 5913 | 3031 | 3317 | 5914 | 3898 | 3049 | 4146 | 3704 | 3779 |
| **DNA-Seq** | **Single ended** | **47** | 1492 | 1212 | 499 | 752 | 868 | 474 | 1010 | 1127 | 971 | 829 | 1052 | 592 | 757 | 745 | 551 |
| | | **90** | 3109 | 1503 | 1156 | 1703 | 1200 | 1105 | 2246 | 1580 | 1299 | 1622 | 1474 | 1135 | 1653 | 1447 | 1509 |
| | | **136** | 4771 | 2436 | 1944 | 2563 | 2111 | 1898 | 3503 | 2878 | 2890 | 2332 | 1948 | 2463 | 1774 | 2715 | 2378 |
| | | **177** | 6572 | 3040 | 2554 | 3566 | 2856 | 2569 | 4497 | 3681 | 3721 | 3672 | 2941 | 3204 | 3436 | 2922 | 3117 |
| | | **245** | 8888 | 4376 | 3800 | 5694 | 4021 | 3638 | 6776 | 5041 | 4526 | 5442 | 4473 | 3558 | 4717 | 3990 | 4131 |
| | **Paired ended** | **47** | 2587 | 1690 | 1008 | 2067 | 1277 | 1020 | 1568 | 1228 | 1125 | 1348 | 1574 | 1274 | 1446 | 1176 | 742 |
| | | **90** | 4413 | 2483 | 1521 | 3628 | 2382 | 1457 | 3477 | 2825 | 2701 | 2692 | 1715 | 2458 | 2432 | 2402 | 2434 |
| | | **136** | 6231 | 3535 | 2331 | 4704 | 3518 | 3118 | 5494 | 4307 | 2902 | 5139 | 3700 | 2513 | 4723 | 3686 | 2477 |
| | | **177** | 8583 | 4520 | 4092 | 6345 | 4607 | 4104 | 6084 | 5483 | 3697 | 5945 | 4620 | 3171 | 5630 | 4170 | 4328 |
| | | **245** | 11533 | 5666 | 6194 | 10091 | 7214 | 6204 | 7227 | 8706 | 7302 | 7646 | 5763 | 6082 | 6907 | 6496 | 4781 |

**Table 3. Duration of compression by ABRIDGE with different parameters and with different compressors**

| | | | Time (in seconds) | | | | | | | | | | | | | | |
|---|---|---|---|---|---|---|---|---|---|---|---|---|---|---|---|---|---|
| | | **Numer of reads (in millions)** | **Parameters Setting 1** | | | **Parameters Setting 2** | | | **Parameters Setting 3** | | | **Parameters Setting 4** | | | **Parameters Setting 5** | | |
| | | | **BR** | **7Z** | **ZPAQ** | **BR** | **7Z** | **ZPAQ** | **BR** | **7Z** | **ZPAQ** | **BR** | **7Z** | **ZPAQ** | **BR** | **7Z** | **ZPAQ** |
| **RNA-Seq** | **Single ended** | **24** | 73 | 103 | 1169 | 81 | 85 | 301 | 57 | 63 | 255 | 58 | 53 | 216 | 45 | 46 | 81 |
| | | **50** | 150 | 211 | 2414 | 165 | 175 | 594 | 114 | 124 | 499 | 97 | 110 | 457 | 89 | 92 | 163 |
| | | **76** | 184 | 271 | 3600 | 202 | 227 | 864 | 146 | 167 | 718 | 130 | 130 | 643 | 110 | 113 | 233 |
| | | **98** | 270 | 353 | 4944 | 290 | 282 | 1203 | 182 | 226 | 1023 | 181 | 187 | 843 | 161 | 148 | 297 |
| | | **128** | 362 | 470 | 6434 | 352 | 392 | 1456 | 272 | 293 | 1314 | 230 | 225 | 1030 | 207 | 194 | 327 |
| | **Paired ended** | **24** | 160 | 188 | 1407 | 249 | 265 | 671 | 236 | 249 | 635 | 220 | 231 | 603 | 195 | 199 | 199 |
| | | **50** | 364 | 432 | 3234 | 530 | 403 | 1623 | 338 | 515 | 1553 | 457 | 337 | 1470 | 264 | 411 | 557 |
| | | **76** | 782 | 629 | 4867 | 817 | 874 | 2293 | 730 | 522 | 2246 | 676 | 712 | 1792 | 604 | 397 | 824 |
| | | **98** | 1004 | 817 | 5767 | 1014 | 1560 | 3569 | 626 | 991 | 2544 | 872 | 609 | 2357 | 775 | 514 | 744 |
| | | **128** | 912 | 1076 | 8257 | 1984 | 1456 | 3910 | 835 | 1295 | 3889 | 763 | 826 | 3645 | 998 | 1024 | 972 |
| **DNA-Seq** | **Single ended** | **47** | 165 | 221 | 2760 | 204 | 201 | 576 | 135 | 140 | 495 | 115 | 122 | 397 | 124 | 129 | 317 |
| | | **90** | 319 | 406 | 4922 | 373 | 394 | 957 | 258 | 277 | 881 | 232 | 238 | 768 | 220 | 233 | 557 |
| | | **136** | 506 | 615 | 7545 | 586 | 627 | 1533 | 386 | 409 | 1214 | 339 | 363 | 1055 | 338 | 365 | 889 |
| | | **177** | 713 | 791 | 10943 | 724 | 812 | 1853 | 507 | 556 | 1560 | 442 | 464 | 1434 | 419 | 460 | 1034 |
| | | **245** | 969 | 1329 | 12875 | 1126 | 1399 | 2702 | 680 | 746 | 2432 | 728 | 632 | 2030 | 577 | 710 | 1476 |
| | **Paired ended** | **47** | 512 | 747 | 3541 | 523 | 543 | 1253 | 472 | 677 | 1131 | 429 | 437 | 814 | 418 | 427 | 661 |
| | | **90** | 1307 | 1267 | 6091 | 1288 | 1283 | 1894 | 1249 | 1050 | 2114 | 939 | 1201 | 1942 | 1157 | 875 | 1273 |
| | | **136** | 2027 | 1676 | 8383 | 1620 | 1693 | 3536 | 1381 | 1454 | 2507 | 1275 | 1340 | 2928 | 1258 | 1267 | 1861 |
| | | **177** | 1987 | 2189 | 10548 | 2686 | 2727 | 4519 | 2425 | 2479 | 3225 | 2279 | 1678 | 3758 | 1590 | 1638 | 3157 |
| | | **245** | 2716 | 3753 | 14544 | 2965 | 3178 | 5198 | 2477 | 2519 | 4636 | 2335 | 2399 | 4107 | 2319 | 2279 | 4234 |

**Table 4. Duration of decompression by ABRIDGE with different parameters and with different compressors**

| | | | Duration (in seconds) | | | | Memory (in MB) | | |
|---|---|---|---|---|---|---|---|---|---|
| | | **Numer of reads (in millions)** | **ABRIDGE** | **BAM** | **CRAM** | | **ABRIDGE** | **BAM** | **CRAM** |
| **RNA-Seq** | **Single ended** | **24** | 9 | 11 | 2 | | 44.44 | 3.5 | 2.82 |
| | | **50** | 9 | 12 | 1 | | 36.14 | 6.05 | 2.84 |
| | | **76** | 18 | 22 | 1 | | 58.65 | 9.28 | 2.62 |
| | | **98** | 24 | 32 | 3 | | 86.17 | 14.11 | 2.77 |
| | | **128** | 31 | 40 | 3 | | 101.09 | 15.4 | 2.79 |
| | **Paired ended** | **24** | 48 | 53 | 6 | | 125.96 | 18.62 | 2.85 |
| | | **50** | 23 | 25 | 2 | | 65.47 | 4.6 | 2.64 |
| | | **76** | 30 | 37 | 4 | | 89.61 | 5.02 | 2.95 |
| | | **98** | 35 | 48 | 4 | | 109.32 | 7.37 | 3 |
| | | **128** | 53 | 58 | 4 | | 126.96 | 8.1 | 3.01 |
| **DNA-Seq** | **Single ended** | **47** | 44 | 22 | 2 | | 48.04 | 2.91 | 2.66 |
| | | **90** | 46 | 21 | 1 | | 42.86 | 2.93 | 2.88 |
| | | **136** | 59 | 40 | 2 | | 64.24 | 4.85 | 2.96 |
| | | **177** | 118 | 60 | 2 | | 86.12 | 2.89 | 3.07 |
| | | **245** | 125 | 75 | 5 | | 105.39 | 3.23 | 3.14 |
| | **Paired ended** | **47** | 150 | 105 | 5 | | 132.4 | 3.14 | 3.26 |
| | | **90** | 89 | 39 | 2 | | 77.77 | 3.09 | 2.84 |
| | | **136** | 110 | 64 | 4 | | 114.85 | 3.02 | 2.85 |
| | | **177** | 139 | 81 | 5 | | 144.95 | 3.01 | 3 |
| | | **245** | 198 | 109 | 6 | | 195.78 | 3.05 | 2.99 |

**Table 5. Duration and memory consumption of ABRIDGE to create index for random search**

| | Parameter setting 1 | Parameter setting 2 | Parameter setting 3 | Parameter setting 4 | Parameter setting 5 |
|---|---|---|---|---|---|
| Save Exact Quality scores | **Yes** | No | No | No | No |
| Save Quality scores | **Yes** | **Yes** | **No** | No | No |
| Save Soft Clippings | **Yes** | **Yes** | **Yes** | No | No |
| Save mismatches | **Yes** | **Yes** | **Yes** | No | No |
| Save unmapped reads | **Yes** | **Yes** | **Yes** | **Yes** | No |

**Table 6. Illustration of arguments provided to ABRIDGE for each parameter setting**

| | | | Memory (in GB) | | | | | |
|---|---|---|---|---|---|---|---|---|
| | | **Numer of reads (in millions)** | **ABRIDGE** | **CSAM** | **DEEZ** | **GENOZIP** | **BAM** | **CRAM** |
| **RNA-Seq** | **Single ended** | **24** | 10.6035538 | 0.01319885 | 3.03070831 | 0.12991714 | 0.00118256 | 0.00125885 |
| | | **50** | 12.1946945 | 0.01726913 | 3.05444717 | 0.70067596 | 0.00118256 | 0.00125885 |
| | | **76** | 12.9527283 | 0.01953888 | 3.0282135 | 0.74595261 | 0.00118256 | 0.00125885 |
| | | **98** | 13.7239113 | 0.02521133 | 3.03667068 | 0.75827408 | 0.00118256 | 0.00125885 |
| | | **128** | 14.7027435 | 0.02589035 | 3.07320023 | 0.79193115 | 0.00118256 | 0.00125885 |
| | **Paired ended** | **24** | 10.4386711 | 0.01242828 | 2.9929924 | 0.13118362 | 0.00118256 | 0.00125885 |
| | | **50** | 10.4650269 | 0.01849747 | 2.99604034 | 0.17809296 | 0.00118256 | 0.00125885 |
| | | **76** | 10.4858246 | 0.01922226 | 3.02975845 | 0.21195984 | 0.00118256 | 0.00125885 |
| | | **98** | 10.5088577 | 0.02093506 | 3.02715302 | 0.25325394 | 0.00118256 | 0.00125885 |
| | | **128** | 10.5120354 | 0.02730179 | 3.04006195 | 0.29411316 | 0.00118256 | 0.00125885 |
| **DNA-Seq** | **Single ended** | **47** | 10.5367737 | 0.03121567 | 3.00671387 | 0.28727341 | 0.00118256 | 0.00125885 |
| | | **90** | 10.6379623 | 0.04970932 | 2.98199844 | 1.22003937 | 0.00118256 | 0.00125885 |
| | | **136** | 11.5782166 | 0.05479431 | 3.00313568 | 1.34737396 | 0.00118256 | 0.00125885 |
| | | **177** | 11.9462471 | 0.08978271 | 3.02417374 | 1.38080978 | 0.00118256 | 0.00125885 |
| | | **245** | 12.5375938 | 0.09642029 | 3.02261353 | 1.54094696 | 0.00118256 | 0.00125885 |
| | **Paired ended** | **47** | 10.5375633 | 0.03024292 | 2.9940834 | 0.26768112 | 0.00118256 | 0.00125885 |
| | | **90** | 10.636692 | 0.05019379 | 3.00325394 | 0.47337723 | 0.00118256 | 0.00125885 |
| | | **136** | 11.5744591 | 0.05505371 | 3.00336456 | 0.67036819 | 0.00118256 | 0.00125885 |
| | | **177** | 11.9455643 | 0.08964157 | 3.02405548 | 0.8114357 | 0.00118256 | 0.00125885 |
| | | **245** | 12.5394821 | 0.09854507 | 3.02558899 | 1.05052567 | 0.00118256 | 0.00125885 |

**Table 7. Memory consumed by different software while accessing a random location**

| | | | Time of access (in seconds) | | | | | |
|---|---|---|---|---|---|---|---|---|
| | | **Numer of reads (in millions)** | **ABRIDGE** | **CSAM** | **DEEZ** | **GENOZIP** | **BAM** | **CRAM** |
| **RNA-Seq** | **Single ended** | **24** | 48 | 1 | 435 | 3 | 0 | 0 |
| | | **50** | 91 | 1 | 920 | 7 | 0 | 0 |
| | | **76** | 131 | 3 | 1421 | 9 | 0 | 0 |
| | | **98** | 168 | 2 | 1884 | 9 | 0 | 0 |
| | | **128** | 215 | 3 | 2476 | 10 | 0 | 0 |
| | **Paired ended** | **24** | 54 | 2 | 432 | 2 | 0 | 0 |
| | | **50** | 100 | 4 | 942 | 3 | 0 | 0 |
| | | **76** | 145 | 3 | 1438 | 4 | 0 | 0 |
| | | **98** | 186 | 3 | 1819 | 4 | 0 | 0 |
| | | **128** | 240 | 7 | 2375 | 5 | 0 | 0 |
| **DNA-Seq** | **Single ended** | **47** | 69 | 2 | 1734 | 5 | 0 | 0 |
| | | **90** | 120 | 1 | 2836 | 35 | 0 | 0 |
| | | **136** | 179 | 7 | 3840 | 36 | 0 | 0 |
| | | **177** | 228 | 25 | 4695 | 41 | 0 | 0 |
| | | **245** | 312 | 3 | 5964 | 42 | 0 | 0 |
| | **Paired ended** | **47** | 77 | 2 | 1688 | 3 | 0 | 0 |
| | | **90** | 135 | 37 | 2706 | 9 | 0 | 0 |
| | | **136** | 203 | 6 | 3699 | 14 | 0 | 0 |
| | | **177** | 262 | 2 | 4459 | 16 | 0 | 0 |
| | | **245** | 357 | 7 | 5846 | 22 | 0 | 0 |

**Table 8. Duration of different software to access a random location**
